# Ultra-high throughput multiplexing and sequencing of >500 bp amplicon regions on the Illumina HiSeq 2500 platform

**DOI:** 10.1101/417618

**Authors:** Johanna B. Holm, Michael S. Humphrys, Courtney K. Robinson, Matthew L. Settles, Sandra Ott, Li Fu, Hongqiu Yang, Pawel Gajer, Xin He, Elias McComb, Patti E Gravitt, Khalil G. Ghanem, Rebecca M. Brotman, Jacques Ravel

## Abstract

Amplification, sequencing and analysis of the 16S rRNA gene affords characterization of microbial community composition. As this tool has become more popular and amplicon-sequencing applications have grown in the total number of samples, growth in sample multiplexing is becoming necessary while maintaining high sequence quality and sequencing depth. Here, modifications to the Illumina HiSeq 2500 platform are described which produce greater multiplexing capabilities and 300 bp paired-end reads of higher quality than produced by the current Illumina MiSeq platform. To improve the feasibility and flexibility of this method, a 2-Step PCR amplification protocol is also described that allows for targeting of different amplicon regions, thus improving amplification success from low bacterial bioburden samples.

**Importance:** Amplicon sequencing has become a popular and widespread tool for surveying microbial communities. Lower overall costs associated with high throughput sequencing have made it a widely-adopted approach, especially for projects which necessitate sample multiplexing to eliminate batch effect and reduced time to acquire data. The method for amplicon sequencing on the Illumina HiSeq 2500 platform described here provides improved multiplexing capabilities while simultaneously producing greater quality sequence data and lower per sample cost relative to the Illumina MiSeq platform, without sacrificing amplicon length. To make this method more flexible to various amplicon targeted regions as well as improve amplification from low biomass samples, we also present and validate a 2-Step PCR library preparation method.

## Introduction

The introduction of the Illumina HiSeq and MiSeq platforms has allowed for the characterization of microbial community composition and structure by enabling in-depth, paired-end sequencing of amplified fragments of the 16S rRNA gene, the ITS region, and other marker genes. The Illumina MiSeq instrument produces paired sequence reads up to 300 bp long. However, low amplicon sequence diversity often results in reduced sequence read quality because of the homogenous signals generated across the entire flow cell [1]. The co-sequencing of PhiX DNA can alleviate the problem, but reduces the overall sequence read throughput and multiplexing options. Alternatively, the addition of a “heterogeneity spacer” in the amplification primer offsets the sequence reads by up to 7 bases and simultaneously increases multiplexing capacity by lowering the amount of PhiX control DNA to ~5% [1]. Lower overall costs associated with high throughput sequencing have made it a widely-adopted approach, especially for projects which necessitate sample multiplexing to eliminate batch effect and reduced time to acquire data, which is often the case in sequencing cores. The Illumina HiSeq 2500 platform with its high throughput offers a remedy to limitations in multiplexing but can currently only be used on short amplicons (i.e. the 16S rRNA gene V4 region) due to limitations in read length (maximum of 250 bp PE in Rapid Run Mode on a HiSeq 2500 instrument) [2].

We present a method that produces high-quality 300 bp paired-end reads (median Q-score 37.1) from up to 1,568 samples per lane on a HiSeq 2500 instrument set to Rapid Run Mode. To make this method feasible and flexible in sequencing different amplicon regions, libraries are prepared using a modified version of previously published 1-Step PCR [1] and 2-Step PCR (https://support.illumina.com/downloads/16s_metagenomic_sequencing_library_preparation.html) methods. In the 1-Step PCR method, fusion primers that contain both the target amplification primer, the heterogeneity spacer, the barcode, and the sequencing primers have been used to amplify a ready-to-sequence amplicon. However, primers ranging from 90-97 bp in length are expensive, can be subject to degradation, and are associated with poor or no amplification from low biomass samples, and are limited to the targeted amplicon region. The 2-Step PCR library preparation procedure described here is more flexible and improves amplification from low biomass samples because the 1^st^ step primers are short, target the amplicon region of interest, and contain the heterogeneity spacer and Illumina Sequencing Primer. The barcodes and flow-cell linker sequences are introduced in a second round of PCR by using the Illumina Sequencing primer as a target.

A previously published 2-Step PCR method [2] used triple barcode-indexing, produced 2 × 250 bp paired-end reads on the Illumina HiSeq 2500 platform, and reported a taxon-specific sequencing bias of the first step primers which differed in both barcode sequence and heterogeneity spacer length. The method we present here uses 8 bp dual-indexing as described by Fadrosh *et al.* [1] wherein the forward index is never used as a reverse index, produces 2 × 300 bp paired end reads by modifying the Illumina HiSeq 2500 sequencing method, and attempts to control for amplification biases by implementing 1) an equimolar ratio of all PCR step 1 primers (which differ only in the length of the heterogeneity spacers) provided to each sample to reduce biases imposed by the heterogeneity spacer, and 2) introduction of barcode sequences are in the second PCR step for library preparation.

In addition to the benefit of flexibility in choice of gene target, we show that the 2-Step PCR method improves amplification success of low biomass samples relative to the 1-Step PCR method. Additionally, we show that the 2-Step PCR method does not significantly bias the measured microbial community by comparing vaginal community state types [3] as defined by taxonomic profiling of vaginal samples of pre- and post-menopausal women [4] targeting the V3-V4 region of the 16S rRNA gene. Post-menopausal vaginal samples tend to be lower in absolute bacterial load relative to pre-menopausal samples [5, 6], making amplification challenging. Samples from each woman were prepared using the 1-Step PCR procedure [1] sequenced on the Illumina MiSeq platform, and the 2-Step PCR procedure sequenced on both the Illumina MiSeq and HiSeq platforms. In addition to comparing the quality of libraries sequenced on the Illumina HiSeq and MiSeq platforms, we also sought to measure 1) improved amplification efficiency of samples prepared by the 2-Step PCR method compared to the 1-Step method and 2) the differences in intra-individual vaginal community state types between methods. Finally, we demonstrate the precision of this method using a comparative mock community analysis.

## Materials & Methods

### Overall Study Design

First, to determine if the choice of library preparation method improved amplification of low-biomass samples, we specifically processed 92 vaginal samples using the dual-indexing 1-Step [1] and 2-Step (described below) library preparation methods. The success of amplifying the 16S rRNA V3V4 region from genomic DNA was evaluated for each method.

To then determine if the choice of library preparation method or sequencing platform impacted the observed sample microbial composition of these samples, we sequenced the libraries of samples successfully produced using both 1-Step and 2-Step methods on the Illumina MiSeq (1-Step and 2-Step) and HiSeq (2-Step only) platforms. The same 2-Step library was sequenced on the MiSeq and HiSeq platforms. The compositions of samples for which high-quality data were obtained from all three methods were statistically compared.

To further validate if sequencing platform impacted observed microbial compositions, we also produced ten separate V3-V4 16S rRNA gene amplicon libraries from the ZymoBIOMICS Microbial Community DNA Standard (Zymo Research, Irvine, CA) using the 2-Step library preparation method, and sequenced each library on separate runs of the Illumina HiSeq platform. We compared the microbial compositions of these samples to theoretical values reported by Zymo as well as to V3-V4 amplicon libraries of the same standard prepared and sequenced by Zymo Research on the Illumina MiSeq platform (see **Supplementary File 7** for library preparation and sequencing methods).

Finally, to compare the sequencing quality and per sample read statistics (per sample number and quality of reads) produced by the Illumina MiSeq and HiSeq 2500 platforms, amplicon libraries from vaginal samples were produced using the 2-Step PCR method and sequenced on both the Illumina MiSeq (276 out of possible 576 samples) and HiSeq 2500 (1,194 out of possible 1,568 samples) platforms. All amplicon libraries targeted the 16S rRNA gene V3-V4 regions from human vaginal samples.

### Genomic DNA extraction

Clinician-collected mid-vaginal ESwabs were stored in Amies transport medium (Copan, Murrieta, CA) as previously described [4]. The study was approved by the University of Maryland Baltimore and the Johns Hopkins School of Public Health Institutional Review Board. Samples were thawed on ice and vortexed briefly. A 0.5 mL aliquot of the cell suspension was transferred to a FastPrep Lysing Matrix B (MP Biomedicals, Santa Ana, CA) tube containing 0.5 mL of PBS (Invitrogen, Carlsbad, CA). A cell lysis solution containing 5 μL lysozyme (10 mg/ml; EMD Chemicals, Gibbstown, NJ), 13 μL mutanolysin (11,700 U/ml; Sigma Aldrich, St. Louis, MO), and 3.2 μL lysostaphin (1 mg/ml; Ambi Products, LLC, Lawrence, NY) was added and samples were incubated at 37°C for 30 min. Then, 10 μL Proteinase K (20mg/ml; Invitrogen), 50 μL 10% SDS (Sigma Aldrich, St. Louis, MO), and 2 μL RNase A (10mg/ml; Invitrogen, Carlsbad, CA) were added and samples were incubated at 55°C for an additional 45 min. Cells were lysed by mechanical disruption on a FastPrep homogenizer at 6 m/s for 40 s, and the lysate was centrifuged on a Zymo Spin IV column at 7000 × g for 1 min. (Zymo Research, Irvine, CA). Lysates were further processed on the QIAsymphony platform using the QS DSP Virus/Pathogen Midi Kit (Qiagen, Hilden, GER) according to the manufacturer’s recommendation. DNA quantification was carried out using the Quant-iT PicoGreen dsDNA assay (Invitrogen).

### Sequencing library construction using 1-Step PCR

Sequencing libraries were constructed by amplifying the 16S rRNA gene V3-V4 regions using the 1-Step PCR amplification protocol previously described [1]. Primer sequences ranged from 90-97 bp depending on the length of the heterogeneity spacer (Table 1). Amplification was performed using Phusion Taq Master Mix (1X, ThermoFisher, Waltham, MA) with 3% DMSO, 0.4 μM each primer, and 5 μL of genomic DNA. A standard volume of genomic DNA was used for each library because genomic DNA concentration was not indicative of the number of 16S rRNA gene targets (**Supplementary File 1**). Cycling conditions were as follows: initial denaturation at 98°C for 30 s, 30 cycles of denaturation at 98°C for 15 s, annealing at 58°C for 15 s, and elongation at 72°C for 15 s, followed by a final elongation step at 72°C for 60 s.

**Table 1.**
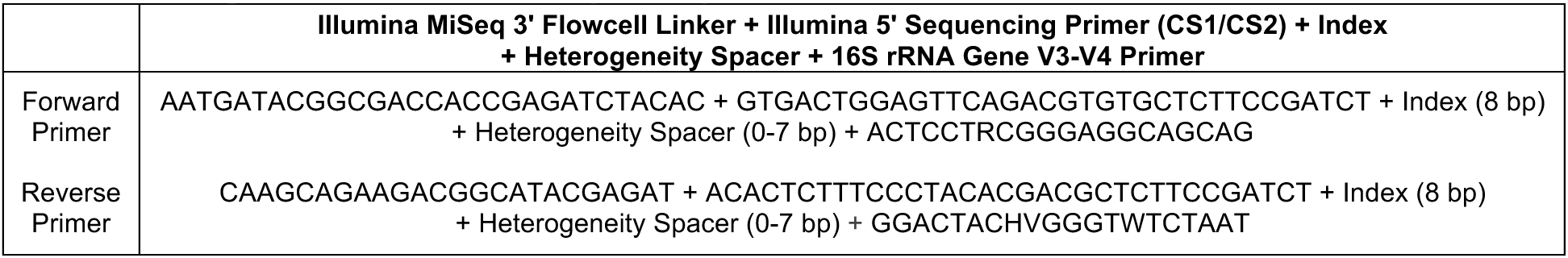
1-Step PCR Method Primers (5’ → 3’)

### Sequencing library construction using 2-Step PCR

The following library preparation method is a modified version of a method provided by Illumina (https://support.illumina.com/downloads/16s_metagenomic_sequencing_library_preparation.html)). The V3-V4 regions of 16S rRNA gene were targeted from genomic DNA using primers bacterial 338F and 806R combined with a heterogeneity spacer of 0-7 bp, and the Illumina Sequencing Primers (Table 2, **Step 1**). A single PCR master mix containing an equal ratio of all primers, which vary by the length of the heterogeneity spacer, was used for all samples. This strategy reduces any amplification biases that may be introduced by the differing lengths of the heterogeneity spacers, and is efficient because the primers do not contain barcode indices (Figure 1). Each PCR reaction contained 1X Phusion Taq Master Mix (ThermoFisher), Step 1 Forward and Reverse primers (0.4 μM each, Supplementary Table 1a), 3% DMSO, and 5 μL of genomic DNA. This standard volume of genomic DNA was used for each library because genomic DNA concentration was not indicative of the number of 16S rRNA gene targets (**Supplementary File 1**). PCR amplification was performed using the following cycling conditions: an initial denaturation at 94°C for 3 min, 20 cycles of denaturation at 94°C for 30 s, annealing at 58°C for 30 s, and elongation at 72°C for 1 min, and a final elongation step at 72°C for 7 min. We used only 20 PCR cycles because biases in microbial community profiles have been reported with higher number of cycles [2]. The resultant amplicons were diluted 1:20, and 1 μL was used in the second step PCR. This second amplification step introduces an 8 bp dual-index barcode to the 16S rRNA gene V3-V4 regions amplicons (Supplementary Table 1b), as well as the flow cell linker adaptors using primers containing a sequence that anneals to the Illumina sequencing primer sequence introduced in step 1 (Table 2, **Step 2** and for full oligonucleotide sequences see **Supplementary Tables 1c and 1d**). Each primer was added to a final concentration of 0.4 μM in each sample specific reaction, along with Phusion Taq Master Mix (1X) and 3% DMSO. Phusion Taq Polymerase (ThermoFisher) was used with the following cycling conditions: an initial denaturation at 94°C for 30 s, 10 cycles consisting of denaturation at 94°C for 30 s, annealing at 58°C for 30 s, and elongation at 72°C for 60 s, followed by a final elongation step at 72°C for 5 min (Figure 1).

**Table 2.**
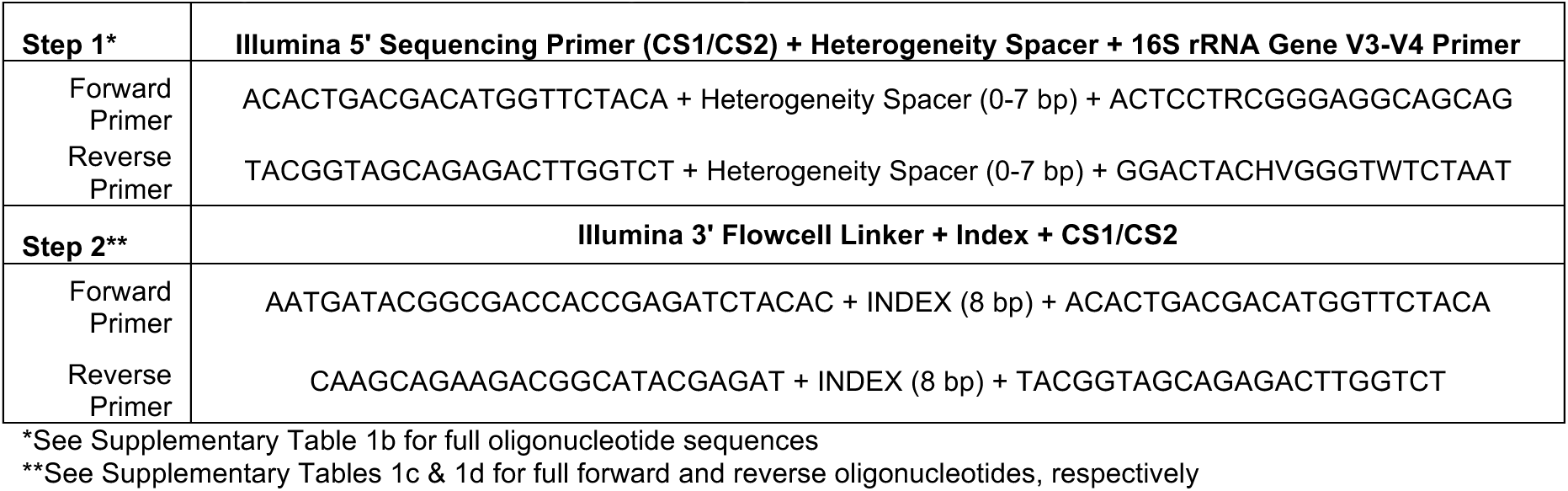
2-Step Protocol PCR Primers (5’ → 3’)

**Figure 1.**
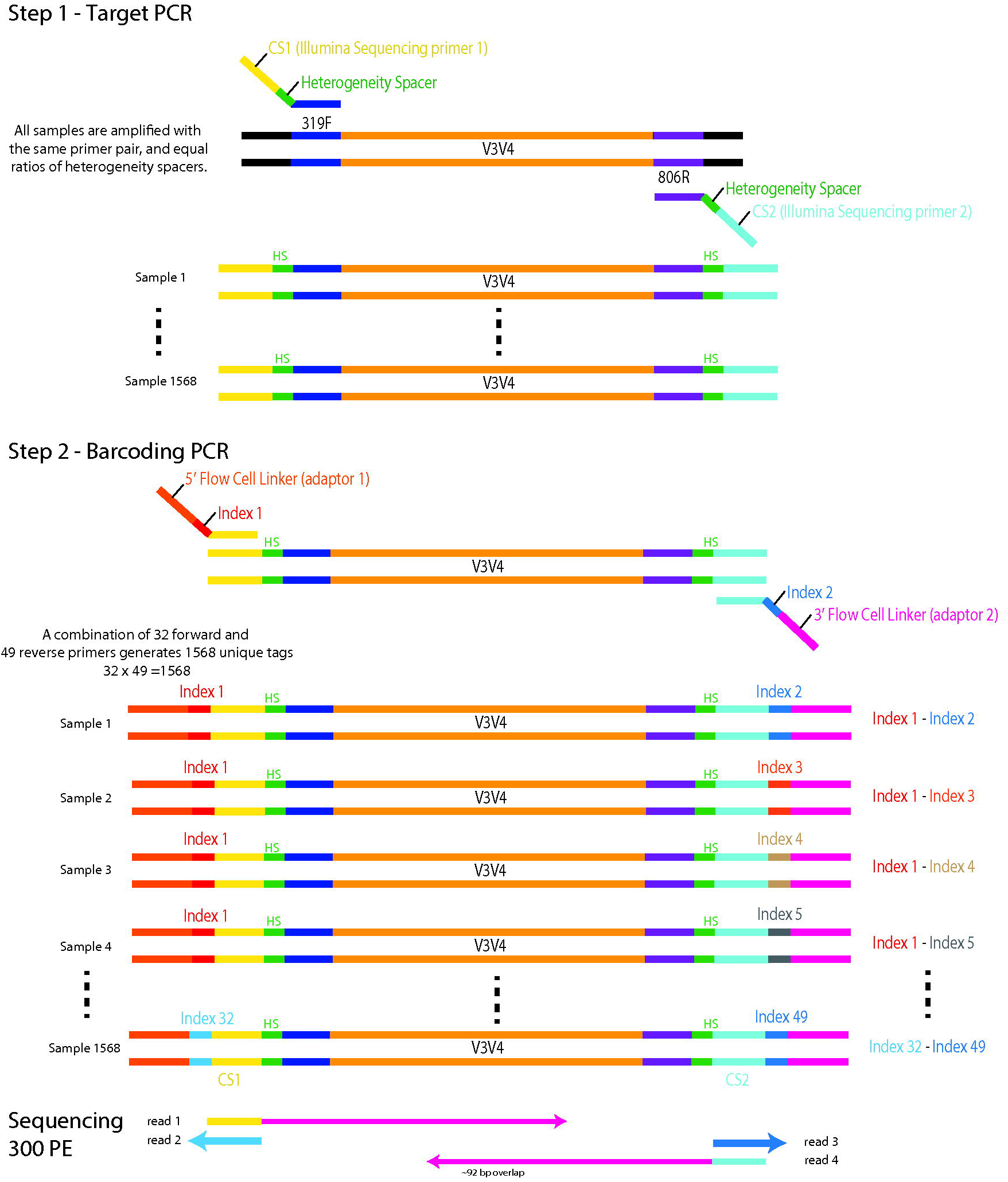
Illumina amplicon library preparation through 2-Step PCR amplification. In the Step 1 PCR, the target gene is amplified using primers that contain the heterogeneity space, and the CS1 lllumina Sequencing primers. The 2nd PCR targets the CS1 lllumina Sequencing primer to add the indices and the illumina flow-cell linker sequence. Sequencing proceeds wherein Reads 1 and 4 contain the forward and reverse target gene sequence, respectively, and Reads 2 and 3 contain the first and second barcode indices, respectively.

### Amplicon library pooling for sequencing

For a large number of samples, library purification and quantification for each sample would be time and labor-intensive. To streamline the process, we visualize libraries on 2% agarose E-Gel (ThermoFisher) and determine the relative amplification success at the expected ~627 bp band size (amplicon + spacer + all primer sequences + linker). Strong, clear bands indicate successful amplification, a weak or fuzzy band indicates intermediate and no band indicates low amplification success. We then standardize the volume of each sample to pool to either 5, 10, or 15 µL of each sample depending on the high, intermediate, or low amplification success of that sample, respectively (see **Supplementary File 2** for labeled gel example). The pooled samples were cleaned up with AMPure XP (Agencourt/Beckman Coulter, Brea, CA) beads following manufacturer’s instructions and size selected around 600 bp. After size-selection the DNA was eluted in water. To ensure proper size of PCR product the pooled libraries were run on Agilent TapeStation 2200 with a DNA1000 tape for quality assurance.

Pooled libraries prepared by 1-Step PCR were sequenced on the Illumina MiSeq platform and those prepared by 2-Step PCR were sequenced on both the Illumina HiSeq and MiSeq platforms following the procedures outlined above.

### Amplification success of vaginal samples using the 1-Step and 2-Step PCR library preparation methods

To determine the success or failure of amplifying the 16S rRNA gene V3-V4 regions from vaginal samples which include samples with low absolute bacterial load using the 1-Step or 2-Step protocols, we evaluated the presence or absence of an amplicon band using agarose gel electrophoresis after the final amplification (in the case of the 2-Step protocol, after the 2^nd^ step). Estimates of absolute bacterial abundance in vaginal samples were determined using real-time quantitative PCR as previously described [7]. **Supplementary File 2** contains an example electrophoresis gel labeled with the volume of the library used for pooling. Samples labeled with “20” show no bands, and in this analysis, represent a failure of amplification. All other samples represent successful amplifications.

### Sequencing by Illumina MiSeq and sequence data processing

Libraries were sequenced on an Illumina MiSeq instrument using 600 cycles producing 2 × 300 bp paired-end reads. The sequences were de-multiplexed using the dual-barcode strategy, a mapping file linking barcode to samples and split_libraries.py, a QIIME-dependent script [8]. The resulting forward and reverse fastq files were split by sample using the QIIME-dependent script split_sequence_file_on_sample_ids.py, and primer sequences were removed using TagCleaner (version 0.16) [9]. Further processing followed the DADA2 Workflow for Big Data and DADA2 (v. 1.5.2) (https://benjjneb.github.io/dada2/bigdata.html, [10], **Supplementary File 3**).

### Sequencing by Illumina HiSeq and sequence data processing

Libraries were sequenced on an Illumina HiSeq 2500 using Rapid Run chemistry and a 515 nm laser barcode reader (a required accessory), and loaded at 8 pmol with 20% PhiX library. Paired-end 300 bp reads were obtained using a HiSeq Rapid SBS Kit v2 (2 × 250 bp, 500 cycles kit) combined with a (2 × 50 bp, 100 cycles kit; alternatively, a single 500 bp kit plus 2 × 50 bp kits can be used). In the HiSeq Control Software, under the Run Configuration tab, within the Flow Cell Setup, the Reagent Kit Type was set to “HiSeq Rapid v2”, and the Flow Cell Type to “HiSeq Rapid Flow Cell v2”. Next, within Recipe, the Index Type was set to “Custom”, the Flow Cell Format to Paired End, and the Cycles set to “301”, “8”, “8”, “301”, for Read 1, Index 1, Index 2, and Read 2, respectively (**Supplementary File 4**). Instead of the standard sequencing primers, custom locked nucleic acid primers were used according to the Fluidigm Access Array User Guide Appendices B and C [11] (the Fluidigm system itself not required). These primers are required for sequencing under the modified conditions, so that the CS1 and CS2 regions can be used as primer binding regions (to produce reads 1 and 2, see Figure 1). The sequences were de-multiplexed using the dual-barcode strategy, a mapping file linking barcode to samples (Supplementary Table 1), and split_libraries.py, a QIIME-dependent script [8]. The resulting forward and reverse fastq files were split by sample using the QIIME-dependent script split_sequence_file_on_sample_ids.py, and primer sequences were removed using TagCleaner (version 0.16) [9]. Further processing followed the DADA2 Workflow for Big Data and DADA2 (v. 1.5.2) [10].

### Intra-individual distance-based bacterial community comparisons of vaginal samples

Samples successfully amplified using both library preparation methods were used for comparative analyses. The 1-Step libraries were sequenced on the Illumina MiSeq Platform and the 2-Step libraries were sequenced on both the Illumina MiSeq and HiSeq platforms. Sequences were quality-filtered and assembled as described above. To fairly compare the Illumina HiSeq and MiSeq platforms, lengths of 255 bp and 225 bp were chosen for hard trimming of forward and reverse reads, respectively, because these were the lengths at which median quality scores decreased below 20 for the worst library (see **Supplementary File 5**, specifically the 2-Step MiSeq F and R reads quality decreases dramatically at approximately these lengths, and so the same lengths were applied to all three methods). Individual reads were further truncated at the base where a quality score of 2 was observed and filtered to contain no ambiguous bases. Additionally, the maximum number of expected errors in a read was set to 2. Reads were assembled only if the overlap between forward and reverse reads, which occurs in the conserved region between V3 and V4, was 100% identical. Chimeras for combined runs removed as per the dada2 protocol. A Kruskal-Wallis test was applied to test if differences in the per sample quality scores differed between the three methods (R Package: stats, Function: kruskal.test). For each of the three quality-filtered datasets, amplification sequence variants (ASVs) generated by DADA2 were individually taxonomically classified using the RDP Naïve Bayesian Classifier [12] trained with the SILVA v128 16S rRNA gene sequence database [13]. ASVs of major vaginal taxa were assigned species-level annotations using speciateIT (version 2.0), a novel and rapid per sequence classifier (http://ravel-lab.org/speciateIT), and verified via BLASTn against the NCBI 16S rRNA gene sequence reference database. Read counts for ASVs assigned to the same taxonomy were summed for each sample. To test for differences in the quality scores of samples prepared and sequenced by the different methods, a Kruskal-Wallis Rank Sum test was applied. To determine if library preparation methods influenced microbial community β-diversity, samples were assigned a vaginal community state type as defined by Jensen-Shannon distances and clustering via Ward linkage [3]. Clusters of Jensen-Shannon distances were visualized using t-Stochastic Neighbor Embedding [14] using 5,000 iterations and perplexity set to 30. Agreement of within-subject assigned CSTs between methods was determined using Fleiss’ Kappa statistic κ [15] (R package: irr v 0.84). Here κ = 0 indicates all CST assignments were dissimilar between the libraries, and κ = 1 indicates identical CST assignments. A κ > 0.75 is considered excellent agreement.

### Comparison of mock community microbial compositions between Illumina HiSeq runs

We used the ZymoBIOMICS Microbial Community DNA Standard (Zymo Research) as a mock community for this analysis. To maintain consistency in taxonomic annotations, we used BLAST and the NCBI Reference database to classify each sequence variant in these analyses. Specific single nucleotide variants produced different taxonomic classifications in the following taxa due to truncation to the V3-V4 amplicon regions: *Bacillus subtilis* to *Bacillus mojavensis*, *Listeria monocytogenes* to *Listeria welshimeri*, *Escherichia coli* to *Escherichia fergusonii*, however, we manually verified the identity of the sequence variants. To determine if the mock community amplicon library compositions produced by the our 2-Step library preparation and HiSeq sequencing methods were within the same variation observed by Zymo Research, we statistically compared the distributions of Jensen-Shannon distances between the Zymo-MiSeq samples and the reported theoretical values and our HiSeq samples and the theoretical values using a Mann-Whitney-Wilcoxon test (R Package: stats, Functions: wilcox.test).

### Sequencing Quality Comparisons of Illumina HiSeq and Illumina MiSeq Sequencing of 2-Step PCR Amplicon Libraries

To compare the sequence quality produced on the near-full Illumina MiSeq and HiSeq runs, the per cycle mean, median, and 1^st^ and 3^rd^ quartiles were calculated from quality scores of sample-specific forward and reverse fastq files in R version 3.4.4 (2018-03-15) using the qa function of the ShortRead package v 1.36.1 [16], data.table v 1.11.4, and ggplot2 v 3.0.0 [17] (R notebook html available upon request). Because quality scores were not normally distributed, a Mann-Whitney-Wilcoxon test was applied to test if differences in the quality scores per cycle differed between the two sequencing platforms (R Package: stats, Functions: shapiro.test and wilcox.test).

All sequence data are available from NCBI SRA under Accession number SRP159872.

## Results

### 2-Step PCR amplicon library preparation improves amplification success of low biomass vaginal samples

Amplification failure was more common in the 1-Step PCR amplification protocol due to the long primers which degrade over time thus reducing amplification efficiency, especially for low biomass samples (i.e. low absolute bacterial load). **Supplementary File 2** contains an example electrophoresis gel labeled with the volume of the library used for pooling. Samples labeled with “20” show no bands, and in this analysis, represent a failure of amplification. All other samples represent successful amplifications. Of 92 low-biomass vaginal samples (mean subject age 48.9), 54% were successfully amplified using the 1-Step PCR protocol, while the 2-Step protocol produced amplifications from 90% of samples (Table 3). Of 42 vaginal samples that did not amplify by the 1-Step method, 55% were from women over the age of 51, the average age of menopause. Thirty-four of these samples successfully amplified using the 2-Step method, an 80% improvement (Supplementary Table 2). A pan-bacterial qPCR analysis confirmed the significantly lower number of 16S rRNA gene targets in the samples which were amplified by 2-Step PCR but not 1-Step PCR (U= 790.5, p = 0.03, **Supplementary File 6)**. Subjects in this group were also significantly older (U=484, p = 0.001). Amplicons were not observed from 8 samples regardless of protocol type, and 1 sample was successfully amplified using the 1-Step but not the 2-Step procedure.

**Table 3.** Summary of sequencing results for vaginal samples. **Supplementary Figure 6** summarizes the pre-quality filtering per-cycle quality scores.

| Library Preparation Method | **1-Step** | **2-Step** |  |
| --- | --- | --- | --- |
| No. samples attempted to amplify | 92 | 92 |  |
| No. samples successfully amplified | 49 | 83 |  |
| Sequencing Platform | MiSeq | MiSeq | HiSeq |
| % Chimeric Sequences Detected | 0.70 | 3.3 | 3.1 |
| Mean No. Non-chimeric, Assembled Sequences per Sample ± SE | 11,080 ± 1506 | 14,282 ± 483 | 50,514 ± 4427 |
| Median Quality Score per Sample [Q1-Q3] | 36.2 [33.5-37.2]* | 34.9 [29.9-36.3]* | 37.1 [33.0-38.0]* |

\*Significant. Kruskal-Wallis H = 187.85, $p < 2.2 \times 10^{-16}$

### Samples successfully amplified by both 1-Step and 2-Step library preparation methods yield similar sequencing metrics on both Illumina MiSeq and HiSeq platforms

Samples successfully amplified using both library preparation methods (n=49) were used to compare the 1-Step and 2-Step library preparation methods sequenced on the Illumina HiSeq (2-Step only) and MiSeq platforms (1-Step and 2-Step). From each combination of methods, 0.7-3% of sequences were detected as chimeras and removed. This yielded on average 11,080 sequences per sample from the 1-Step library sequenced on the MiSeq platform, 14,282 sequences per sample from the 2-Step library sequenced on the MiSeq platform, and 50,514 sequences per sample from the 2-Step library sequenced on the HiSeq platform (Table 3). Due to low total read counts from some samples, only 30 samples containing > 500 total sequences in each method were used for comparative β-diversity analysis between the three methods. Consistency of observed vaginal community state types (CSTs) between libraries was tested using Fleiss’ *kappa* for inter-rater reliability, where κ > 0.75 indicated excellent agreement. Complete agreement between all three methods was observed and samples clustered primarily by vaginal community state type and subject as opposed to library preparation method or sequencing platform (κ = 1.0, Figure 2, raw read count taxonomy tables are available in Supplementary Table 3).

**Figure 2.**
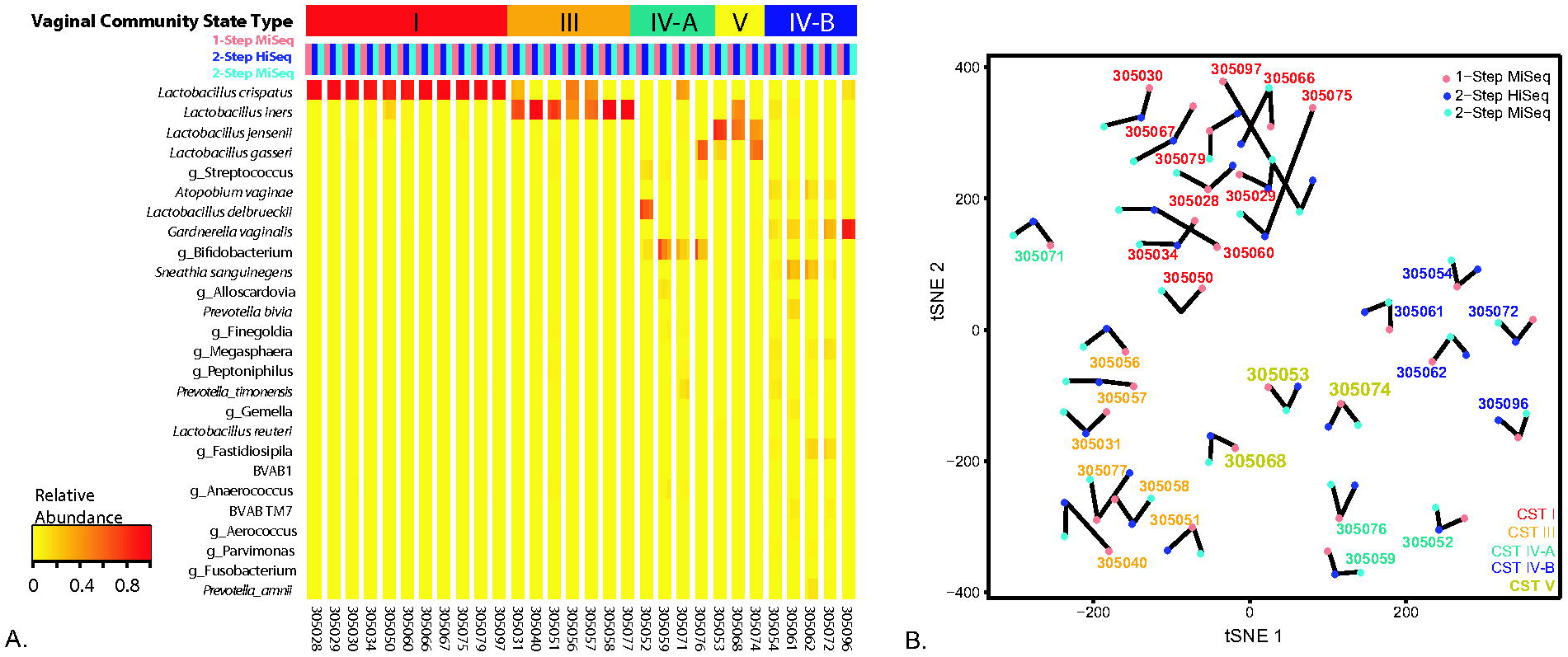
Heatmap of taxon relative abundances (rows) of samples (columns). Subject samples are separate by white lines and samples are ordered by vaginal community state types and as follows: 1-Step MiSeq (pink), 2-step HiSeq (blue), 2-Step MiSeq (aqua). B) tSNE representation of Jensen-Shannon distances between samples from the same subject. Samples primarily cluster by vaginal CST.

### Mock community libraries prepared via 2-Step PCR and sequenced on Illumina HiSeq are not different than those sequenced on Illumina MiSeq

In order to verify that consistency of results was not simply due to sample type, we also compared the microbial compositions of the Zymobiomics Microbial DNA Standard obtained on the Illumina HiSeq and MiSeq platforms. We used theoretical values reported by Zymo, as well as compositional data produced from 16S rRNA gene V3-V4 regions amplicon libraries sequenced on the Illumina MiSeq (prepared, sequenced, and provided by Zymo). The raw read count taxonomy table is available in Supplementary Table 4. The distribution of Jensen-Shannon distances between Zymo-prepared, MiSeq-sequenced microbiota composition and theoretical composition did not significantly differ from the distribution of distances between the 2-Step-prepared, Illumina HiSeq-sequenced and theoretical microbiota compositions (U = 29, p = 0.9578, **Supplementary File 8**).

**Table 4.** Sequencing run information for the MiSeq and HiSeq platforms.

| Sequencing Platform | **MiSeq** | **HiSeq 2500 RR** |
| --- | --- | --- |
| Run Details | 2 x 300 bp PE | 2 x 250 bp<br>+ 2 x 50bp |
| Mean No. Assembled Sequences per Sample ± SE | 13,116 ± 479 | 49,851 ± 895 |
| No. Samples | 276 | 1194 |
| % Chimeric Sequences Detected | 2.8 | 7.7 |
| Mean No. Non-chimeric, Assembled Sequences per Sample ± SE | 12,737 ± 463 | 45,988 ± 787 |
| Median Quality Score [Q25-Q75] – Forward Reads | 35.7 [33-37]* | 36.1 [35-38]* |
| Median Quality Score [Q25-Q75] – Reverse Reads | 33.9 [25-36]<sup>†</sup> | 33.2 [31-37]<sup>†</sup> |
| Cost of Sequencing per Sample (No. Multiplexed Samples) | \$6.38 (384) | \$3.99 (1,568) |

\* Significant. Wilcoxon Rank Sum W = 70352, $p < 2.2 \times 10^{-16}$
<sup>†</sup> Significant. Wilcoxon Rank Sum W = 76453, $p < 2.2 \times 10^{-16}$

### Comparison of Illumina MiSeq and Illumina HiSeq amplicon sequence read quality and quantity

To compare the quality of amplicon sequence reads produced via 2-Step PCR and the Illumina MiSeq and HiSeq platforms, each sequencing run was demultiplexed with the same mapping file, and the sequence reads quality profiles were compared. The two runs had 183 samples in common. Significantly greater mean quality scores of both forward and reverse reads were observed for 1,194 samples run on the HiSeq platform compared to 276 samples run on the MiSeq platform (p < 2.2 × 10^−16^, Figure 3). The HiSeq 2500 platform produced a greater mean number of quality-filtered sequences per sample than the MiSeq platform, with fewer chimeric sequences detected on average (Table 4). Additionally, the HiSeq 2500 sequencing strategy was more cost efficient (nearly 40% cheaper per sample), assuming 2 lanes are run with 1,568 multiplexed samples per lane (Table 4). These results were also consistent across multiple sequencing runs (**Supplementary File 9**).

## Discussion

Microbiome analyses large enough to achieve adequate statistical power are becoming more desirable, and reduced sequencing costs make these analyses feasible. Therefore, ultra-high-throughput sequencing capabilities are needed that do not sacrifice sequence quality. Ideally, such methods would allow for flexibility to target a diverse set of genes or gene regions (for example, ITS regions, the 16S and 23S rRNA genes, the *cpn60* gene [18, 19] among others) while also maintaining the ability to sequence longer amplicons (i.e., the 16S rRNA gene V3-V4 region). The method presented here improves on current technologies by producing consistent high-quality, 300 bp paired-end reads. Relative to the Illumina MiSeq platform, sequencing on the Illumina HiSeq platform produced a greater number of reads per sample, of significantly higher quality, with the capability to multiplex up to 2 × 1,568 samples. The innovative use of the Illumina HiSeq 2500 platform as presented here and by Muinck *et al.* [2] allow for ultra-high-throughput sequencing of amplicon libraries.

**Figure 3.**
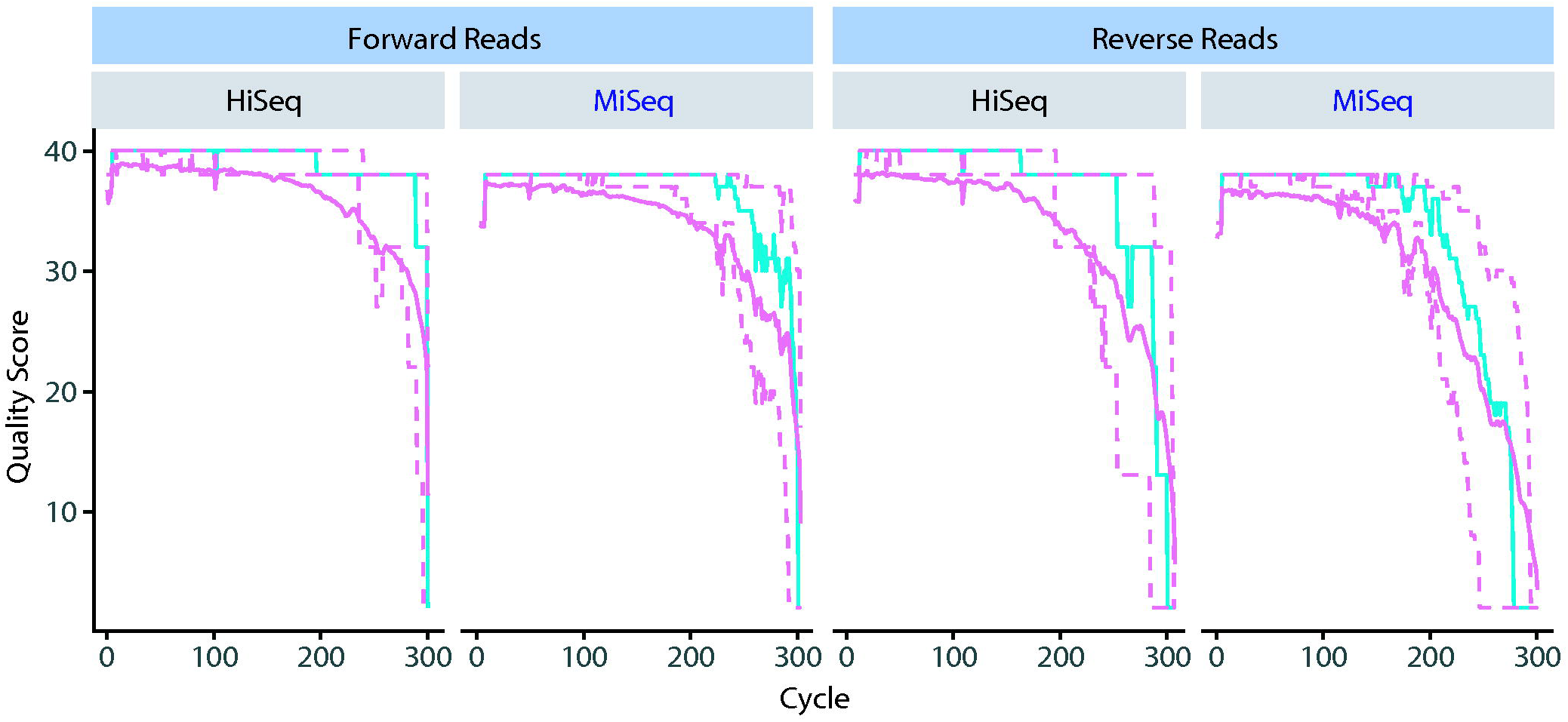
Forward and reverse read quality profiles for 300 cycles on the lllumina HiSeq (1,536 samples and MiSeq (444 samples) platforms. Amplicon libraries were prepared using a 2-Step PCR method. Shown for each cycle are the mean quality score (green line), the median quality score (solid purple line), the quartiles of the quality score distribution (dotted purple lines).

In addition, the 2-Step PCR library preparation method described here makes production of sequencing libraries from various gene targets and samples containing low bacterial loads easy through the use of unindexed, target specific primers in the first round of PCR. Amplification success of samples with low bacterial loads are prone to amplification difficulties, and amplification using the longer primers required in the traditional 1-Step protocol [1] exacerbate the problem because of primer degradation and poor annealing due to the long overhang unprimed sequence. Using the 2-Step PCR approach, we showed an 80% improvement of samples containing low bacterial loads over the 1-Step PCR method. In addition, the shorter primers used in the 2-Step PCR library protocol do not require PAGE purification, lowering the overall cost of the method relative to the 1-Step PCR protocol. Other low-biomass environments that could benefit from this 2-Step PCR procedure include blood and serum [20], respiratory airways [21], skin [22], sub-seafloor sediments [23], and clean rooms [24].

In summary, to demonstrate the comparability of sequence datasets produced via different methods, 16S rRNA gene V3-V4 regions sequence datasets were generated from low-biomass vaginal samples women using both 1-Step and 2-Step PCR library construction methods and the Illumina HiSeq and MiSeq sequencing platforms. Complete within-subject agreement between the vaginal community state type assignments [3] were observed between all three methods, though a greater number of significantly higher quality sequences were obtained from the 2-Step PCR method sequenced on the Illumina HiSeq 2500 platform. We also show that resulting microbial compositions of mock community samples are not significantly altered when amplicon libraries are prepared using the 2-Step library preparation method and sequenced on the Illumina HiSeq platform. We therefore conclude that while the 2-Step PCR preparation method combined with the Illumina HiSeq 2500 platform is preferred, data generated by 1-Step or 2-Step PCR and sequenced on the Illumina MiSeq or HiSeq 2500 platform can be combined to successfully obtain meaningful conclusions about the environment and sample types of interest (given that the same region is targeted).

### Limitations

The method is extremely high-throughput, and as such might not be suitable for small projects unless these are combined with other samples. Producing a large number of samples ready for pooling requires automation so that time from sample collection to data generation is still reasonable. Overall, automation is required, and this approach might be suitable for microbiome service cores where faster turn-around is needed and running many MiSeq runs is not a viable option because of potential batch effect.

## Acknowledgements

JB Holm was supported by the National Institute of Allergy and Infectious Diseases of the National Institutes of Health under award number F32AI136400. Research reported in this publication was supported in part by the National Institute of Allergy and Infectious Diseases and the National Institute of Nursing Research of the National Institutes of Health under award numbers: U19AI084044, R01AI116799, R21AI107224, R01AI089878 and R01NR015495. The content is solely the responsibility of the authors and does not necessarily represent the official views of the National Institutes of Health.

