## Supplemental File 1 for "Ultra-high throughput multiplexing and sequencing of >500 bp amplicon regions on the Illumina HiSeq 2500 platform"

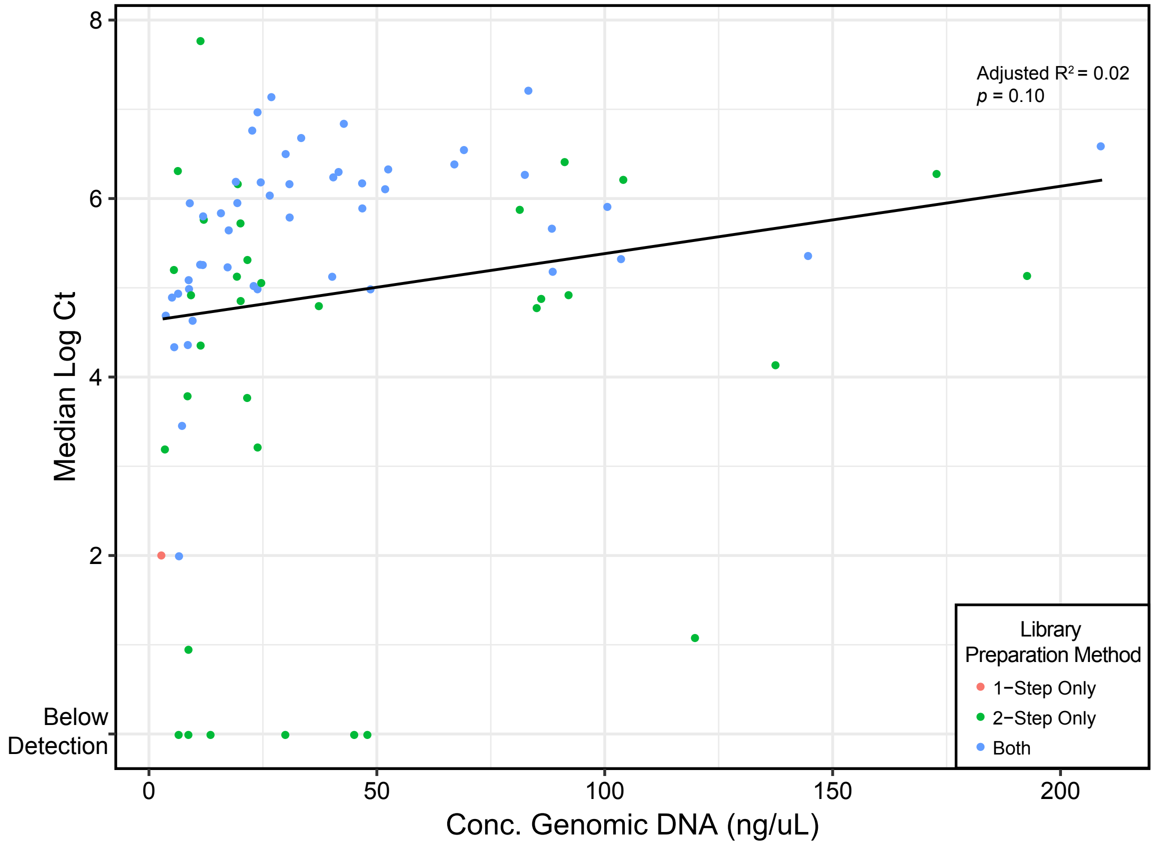


Supplemental File 1. The concentration of genomic DNA extracted from vaginal samples does not correlate to the absolute bacterial load as measured by qPCR of pan-bacterial 16S rRNA genes.
