## Supplemental File 2 for "Ultra-high throughput multiplexing and sequencing of >500 bp amplicon regions on the Illumina HiSeq 2500 platform"

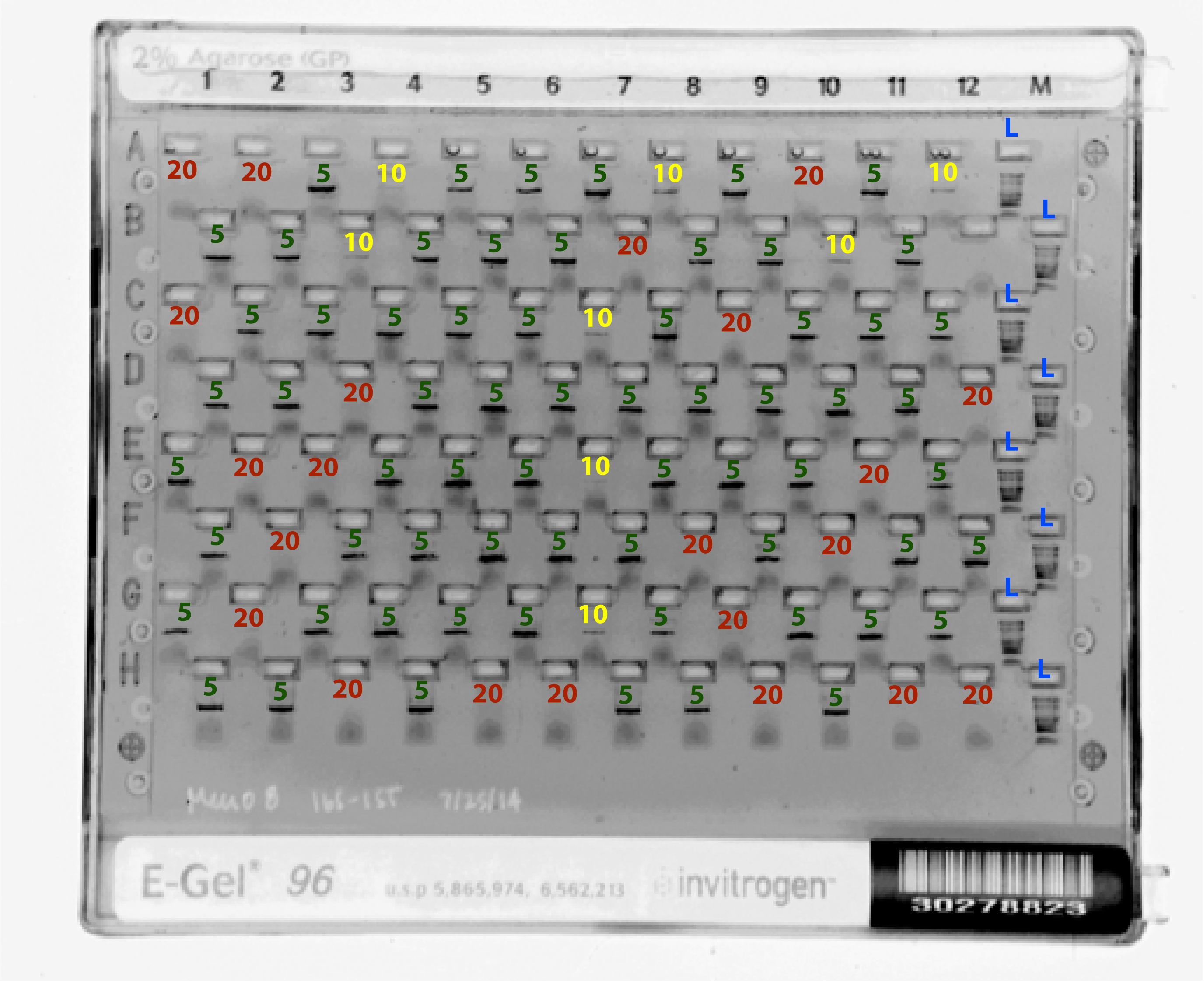


Supplemental File 2. Amplicon libraries are run on a 2% E-Gel to determine amplification succes and the volume of each library to be pooled for sequencing. Strong, crisp bands are successful amplifications for which 5 μL are used in pooling (green “5”s). For samples in which no band is observed, 20 μL are used in pooling (red “20”s). Finally, 10 μL are used in pooling (yellow “10”s) from samples which produce weaker, fuzzy bands.
