## Supplemental File 3 for "Ultra-high throughput multiplexing and sequencing of >500 bp amplicon regions on the Illumina HiSeq 2500 platform"

QIIME1 Code used for sample processing:

QIIME v1.9

NOTE: Some systems label sequencing files as R1, R2, R3, R4, where R1 and R4 are forward and reverse reads, respectively, and R2 and R3 are forward and reverse indices, respectively. 
Other systems label these same files as R1, I1, I2, R2, respectively. The below code accounts for this.

R1 - forward read file
R2/I1 - forward index file
R3/I2 - reverse index file
R4/R2 - reverse read file

extract_barcodes.py --input_type barcode_paired_end -f <raw R2/I1 fastq file> -r <raw R3/I2 fastq file> --bc1_len 8 --bc2_len 8 -o <out directory>
	=> Produces barcodes.fastq file

split_libraries_fastq.py -i <raw R1 fastq file> -o <user-specified R1 out directory> -b barcodes.fastq -m <mapping-file> --max_barcode_errors 1 --store_demultiplexed_fastq --barcode_type 24 -r 999 -n 999 -q 0 -p 0.0001
	=> Produces seqs.fastq file within user-specified R1 out directory

split_libraries_fastq.py -i <raw R4/R2 fastq file> -o <user-specified R4 out directory> -b barcodes.fastq -m <mapping-file> --max_barcode_errors 1 --store_demultiplexed_fastq --barcode_type 24 -r 999 -n 999 -q 0 -p 0.0001
	=> Produces seqs.fastq file within user-specified R4/R2 out directory

split_sequence_file_on_sample_ids.py -i <seqs.fastq from user-specified R1 out directory> --file_type fastq -o <user-specified R1 split files directory>
	=> Produces sample-specific forward read fastq files
split_sequence_file_on_sample_ids.py -i <seqs.fastq from user-specified R4/R2 out directory> --file_type fastq -o <user-specified R4/R2 split files directory>
	=> Produces sample-specific reverse read fastq files

for each sample-specific fastq file:
perl /usr/local/packages/tagcleaner-0.16/bin/tagcleaner.pl -fastq <sample-specific forward read fastq file> -out <fastq-prefix>_R1_tc -line_width 0 -verbose -tag5 GGACTACHVGGGTWTCTAAT -mm5 2 -trim_within 50

perl /usr/local/packages/tagcleaner-0.16/bin/tagcleaner.pl -fastq <sample-specific reverse read fastq file> -out <fastq-prefix>_R2_tc -line_width 0 -verbose -tag5 ACTCCTACGGGAGGCAGCAG -mm5 2 -trim_within 50


DADA2 Code used for sample processing:

R version 3.4.0 (2017-04-21) -- "You Stupid Darkness"
Copyright (C) 2017 The R Foundation for Statistical Computing
Platform: x86_64-pc-linux-gnu (64-bit)
> library("dada2")
Loading required package: Rcpp
> packageVersion("dada2")
[1] ‘1.6.0’
>   ## perform filtering and trimming
>   path <- cwd
>   filtpath <- file.path(path, "filtered")
>   fastqFs <- sort(list.files(path, pattern="R1_tc.fastq"))
>   fastqRs <- sort(list.files(path, pattern="R2_tc.fastq"))
>   sample.names <- sapply(strsplit(basename(fastqFs), "_"), `[`, 1)
>   filtFs<-(paste0(sample.names, "_F_filt.fastq.gz"))
>   filtRs<-(paste0(sample.names, "_R_filt.fastq.gz"))
>   filtFs_files<-file.path(filtpath,filtFs)
>   filtRs_files<-file.path(filtpath,filtRs)
>   if(length(fastqFs) != length(fastqRs)) stop("Forward and reverse files do not match.")
>   out<-filterAndTrim(fwd=file.path(path,fastqFs), filt=filtFs_files, rev=file.path(path,fastqRs), filt.rev=filtRs_files, truncLen=c(255,225), maxN=0, maxEE=c(2,2), truncQ=2, rm.phix=TRUE, compress=TRUE, multithread=TRUE, verbose=TRUE, matchIDs=TRUE)
>
>   ## Learn errors
>   filtFs <- list.files(filtpath, pattern="_F_filt.fastq.gz", full.names = TRUE)
>   filtRs <- list.files(filtpath, pattern="_R_filt.fastq.gz", full.names = TRUE)
>   sample.names <- sapply(strsplit(basename(filtFs), "_"), `[`, 1)
>   sample.namesR <- sapply(strsplit(basename(filtRs), "_"), `[`, 1)
>   if(!identical(sample.names, sample.namesR)) stop("Forward and reverse files do not match.")
>   names(filtFs) <- sample.names
>   names(filtRs) <- sample.namesR
>   set.seed(100)
>   # Learn forward error rates
>   errF <- learnErrors(filtFs, nread=1e6, multithread=TRUE)
Initializing error rates to maximum possible estimate.
Sample 1 - 8442 reads in 1857 unique sequences.
Sample 2 - 18688 reads in 1944 unique sequences.
Sample 3 - 13353 reads in 2789 unique sequences.
Sample 4 - 11985 reads in 1484 unique sequences.
Sample 5 - 21021 reads in 2044 unique sequences.
Sample 6 - 12868 reads in 2019 unique sequences.
Sample 7 - 16892 reads in 2212 unique sequences.
Sample 8 - 20172 reads in 1712 unique sequences.
Sample 9 - 56 reads in 23 unique sequences.
Sample 10 - 21535 reads in 2764 unique sequences.
Sample 11 - 13879 reads in 2216 unique sequences.
Sample 12 - 15009 reads in 2195 unique sequences.
Sample 13 - 3 reads in 3 unique sequences.
Sample 14 - 7 reads in 5 unique sequences.
Sample 15 - 11073 reads in 1934 unique sequences.
Sample 16 - 18565 reads in 1862 unique sequences.
Sample 17 - 18106 reads in 3737 unique sequences.
Sample 18 - 14973 reads in 2160 unique sequences.
Sample 19 - 15268 reads in 2021 unique sequences.
Sample 20 - 6648 reads in 1345 unique sequences.
Sample 21 - 27 reads in 19 unique sequences.
Sample 22 - 19175 reads in 2361 unique sequences.
Sample 23 - 19447 reads in 2527 unique sequences.
Sample 24 - 4605 reads in 1035 unique sequences.
Sample 25 - 17641 reads in 3661 unique sequences.
Sample 26 - 21725 reads in 2411 unique sequences.
Sample 27 - 8890 reads in 1512 unique sequences.
Sample 28 - 12675 reads in 2465 unique sequences.
Sample 29 - 10890 reads in 1914 unique sequences.
Sample 30 - 12911 reads in 4312 unique sequences.
Sample 31 - 19733 reads in 2808 unique sequences.
Sample 32 - 12694 reads in 1639 unique sequences.
Sample 33 - 16816 reads in 1788 unique sequences.
Sample 34 - 16220 reads in 3448 unique sequences.
Sample 35 - 12869 reads in 1881 unique sequences.
Sample 36 - 15883 reads in 4207 unique sequences.
Sample 37 - 17427 reads in 4121 unique sequences.
Sample 38 - 16687 reads in 1888 unique sequences.
Sample 39 - 7962 reads in 1218 unique sequences.
Sample 40 - 14220 reads in 2052 unique sequences.
Sample 41 - 18357 reads in 2142 unique sequences.
Sample 42 - 17902 reads in 1740 unique sequences.
Sample 43 - 15676 reads in 2556 unique sequences.
Sample 44 - 15515 reads in 2991 unique sequences.
Sample 45 - 16520 reads in 3497 unique sequences.
Sample 46 - 14354 reads in 3205 unique sequences.
Sample 47 - 21998 reads in 4487 unique sequences.
Sample 48 - 6805 reads in 2423 unique sequences.
Sample 49 - 12507 reads in 1765 unique sequences.
Sample 50 - 16306 reads in 1730 unique sequences.
Sample 51 - 9980 reads in 1871 unique sequences.
Sample 52 - 13725 reads in 1651 unique sequences.
Sample 53 - 10257 reads in 1460 unique sequences.
Sample 54 - 20581 reads in 2062 unique sequences.
Sample 55 - 16190 reads in 3239 unique sequences.
Sample 56 - 18965 reads in 2763 unique sequences.
Sample 57 - 21372 reads in 5457 unique sequences.
Sample 58 - 14122 reads in 2047 unique sequences.
Sample 59 - 16937 reads in 1864 unique sequences.
Sample 60 - 3329 reads in 1840 unique sequences.
Sample 61 - 18411 reads in 2248 unique sequences.
Sample 62 - 18328 reads in 2261 unique sequences.
Sample 63 - 15051 reads in 1736 unique sequences.
Sample 64 - 22257 reads in 2560 unique sequences.
Sample 65 - 815 reads in 417 unique sequences.
Sample 66 - 18885 reads in 2364 unique sequences.
Sample 67 - 21374 reads in 5466 unique sequences.
Sample 68 - 20498 reads in 2577 unique sequences.
Sample 69 - 2629 reads in 684 unique sequences.
Sample 70 - 3315 reads in 986 unique sequences.
Sample 71 - 13718 reads in 1801 unique sequences.
Sample 72 - 18857 reads in 1639 unique sequences.
   selfConsist step 2 
   selfConsist step 3 
   selfConsist step 4 
Convergence after  4  rounds.
Total reads used:  1012576 
>   # Learn reverse error rates
>   errR <- learnErrors(filtRs, nread=1e6, multithread=TRUE)
Initializing error rates to maximum possible estimate.
Sample 1 - 8442 reads in 1848 unique sequences.
Sample 2 - 18688 reads in 1601 unique sequences.
Sample 3 - 13353 reads in 2455 unique sequences.
Sample 4 - 11985 reads in 1202 unique sequences.
Sample 5 - 21021 reads in 1209 unique sequences.
Sample 6 - 12868 reads in 1264 unique sequences.
Sample 7 - 16892 reads in 1907 unique sequences.
Sample 8 - 20172 reads in 1463 unique sequences.
Sample 9 - 56 reads in 30 unique sequences.
Sample 10 - 21535 reads in 2467 unique sequences.
Sample 11 - 13879 reads in 2667 unique sequences.
Sample 12 - 15009 reads in 2062 unique sequences.
Sample 13 - 3 reads in 3 unique sequences.
Sample 14 - 7 reads in 7 unique sequences.
Sample 15 - 11073 reads in 1766 unique sequences.
Sample 16 - 18565 reads in 1488 unique sequences.
Sample 17 - 18106 reads in 3560 unique sequences.
Sample 18 - 14973 reads in 1908 unique sequences.
Sample 19 - 15268 reads in 1873 unique sequences.
Sample 20 - 6648 reads in 1221 unique sequences.
Sample 21 - 27 reads in 20 unique sequences.
Sample 22 - 19175 reads in 1501 unique sequences.
Sample 23 - 19447 reads in 2419 unique sequences.
Sample 24 - 4605 reads in 1027 unique sequences.
Sample 25 - 17641 reads in 3319 unique sequences.
Sample 26 - 21725 reads in 2095 unique sequences.
Sample 27 - 8890 reads in 1610 unique sequences.
Sample 28 - 12675 reads in 2179 unique sequences.
Sample 29 - 10890 reads in 1661 unique sequences.
Sample 30 - 12911 reads in 3682 unique sequences.
Sample 31 - 19733 reads in 2576 unique sequences.
Sample 32 - 12694 reads in 1436 unique sequences.
Sample 33 - 16816 reads in 1522 unique sequences.
Sample 34 - 16220 reads in 3604 unique sequences.
Sample 35 - 12869 reads in 1508 unique sequences.
Sample 36 - 15883 reads in 3381 unique sequences.
Sample 37 - 17427 reads in 3334 unique sequences.
Sample 38 - 16687 reads in 1873 unique sequences.
Sample 39 - 7962 reads in 1040 unique sequences.
Sample 40 - 14220 reads in 1984 unique sequences.
Sample 41 - 18357 reads in 2153 unique sequences.
Sample 42 - 17902 reads in 1708 unique sequences.
Sample 43 - 15676 reads in 2234 unique sequences.
Sample 44 - 15515 reads in 2469 unique sequences.
Sample 45 - 16520 reads in 3128 unique sequences.
Sample 46 - 14354 reads in 2972 unique sequences.
Sample 47 - 21998 reads in 4250 unique sequences.
Sample 48 - 6805 reads in 2490 unique sequences.
Sample 49 - 12507 reads in 1403 unique sequences.
Sample 50 - 16306 reads in 1639 unique sequences.
Sample 51 - 9980 reads in 1629 unique sequences.
Sample 52 - 13725 reads in 1351 unique sequences.
Sample 53 - 10257 reads in 1895 unique sequences.
Sample 54 - 20581 reads in 2008 unique sequences.
Sample 55 - 16190 reads in 3254 unique sequences.
Sample 56 - 18965 reads in 3327 unique sequences.
Sample 57 - 21372 reads in 4787 unique sequences.
Sample 58 - 14122 reads in 1586 unique sequences.
Sample 59 - 16937 reads in 1318 unique sequences.
Sample 60 - 3329 reads in 1959 unique sequences.
Sample 61 - 18411 reads in 2091 unique sequences.
Sample 62 - 18328 reads in 1707 unique sequences.
Sample 63 - 15051 reads in 1602 unique sequences.
Sample 64 - 22257 reads in 2467 unique sequences.
Sample 65 - 815 reads in 410 unique sequences.
Sample 66 - 18885 reads in 1907 unique sequences.
Sample 67 - 21374 reads in 4237 unique sequences.
Sample 68 - 20498 reads in 2442 unique sequences.
Sample 69 - 2629 reads in 671 unique sequences.
Sample 70 - 3315 reads in 1094 unique sequences.
Sample 71 - 13718 reads in 1964 unique sequences.
Sample 72 - 18857 reads in 1495 unique sequences.
   selfConsist step 2 
   selfConsist step 3 
   selfConsist step 4 
Convergence after  4  rounds.
Total reads used:  1012576 
>   # Sample inference and merger of paired-end reads
>   mergers <- vector("list", length(sample.names))
>   names(mergers) <- sample.names
>   for(sam in sample.names) {
+     cat("Processing:", sam, "
+ ")
+       derepF <- derepFastq(filtFs[[sam]])
+       ddF <- dada(derepF, err=errF, multithread=TRUE)
+       derepR <- derepFastq(filtRs[[sam]])
+       ddR <- dada(derepR, err=errR, multithread=TRUE)
+       merger <- mergePairs(ddF, derepF, ddR, derepR)
+       mergers[[sam]] <- merger
+   }
Processing: 305025 
Sample 1 - 8442 reads in 1857 unique sequences.
Sample 1 - 8442 reads in 1848 unique sequences.
Processing: 305026 
Sample 1 - 18688 reads in 1944 unique sequences.
Sample 1 - 18688 reads in 1601 unique sequences.
Processing: 305027 
Sample 1 - 13353 reads in 2789 unique sequences.
Sample 1 - 13353 reads in 2455 unique sequences.
Processing: 305028 
Sample 1 - 11985 reads in 1484 unique sequences.
Sample 1 - 11985 reads in 1202 unique sequences.
Processing: 305029 
Sample 1 - 21021 reads in 2044 unique sequences.
Sample 1 - 21021 reads in 1209 unique sequences.
Processing: 305030 
Sample 1 - 12868 reads in 2019 unique sequences.
Sample 1 - 12868 reads in 1264 unique sequences.
Processing: 305031 
Sample 1 - 16892 reads in 2212 unique sequences.
Sample 1 - 16892 reads in 1907 unique sequences.
Processing: 305032 
Sample 1 - 20172 reads in 1712 unique sequences.
Sample 1 - 20172 reads in 1463 unique sequences.
Processing: 305033 
Sample 1 - 56 reads in 23 unique sequences.
Sample 1 - 56 reads in 30 unique sequences.
Processing: 305034 
Sample 1 - 21535 reads in 2764 unique sequences.
Sample 1 - 21535 reads in 2467 unique sequences.
Processing: 305035 
Sample 1 - 13879 reads in 2216 unique sequences.
Sample 1 - 13879 reads in 2667 unique sequences.
Processing: 305036 
Sample 1 - 15009 reads in 2195 unique sequences.
Sample 1 - 15009 reads in 2062 unique sequences.
Processing: 305037 
Sample 1 - 3 reads in 3 unique sequences.
Sample 1 - 3 reads in 3 unique sequences.
Processing: 305038 
Sample 1 - 7 reads in 5 unique sequences.
Sample 1 - 7 reads in 7 unique sequences.
Processing: 305039 
Sample 1 - 11073 reads in 1934 unique sequences.
Sample 1 - 11073 reads in 1766 unique sequences.
Processing: 305040 
Sample 1 - 18565 reads in 1862 unique sequences.
Sample 1 - 18565 reads in 1488 unique sequences.
Processing: 305041 
Sample 1 - 18106 reads in 3737 unique sequences.
Sample 1 - 18106 reads in 3560 unique sequences.
Processing: 305042 
Sample 1 - 14973 reads in 2160 unique sequences.
Sample 1 - 14973 reads in 1908 unique sequences.
Processing: 305043 
Sample 1 - 15268 reads in 2021 unique sequences.
Sample 1 - 15268 reads in 1873 unique sequences.
Processing: 305044 
Sample 1 - 6648 reads in 1345 unique sequences.
Sample 1 - 6648 reads in 1221 unique sequences.
Processing: 305045 
Sample 1 - 27 reads in 19 unique sequences.
Sample 1 - 27 reads in 20 unique sequences.
Processing: 305046 
Sample 1 - 19175 reads in 2361 unique sequences.
Sample 1 - 19175 reads in 1501 unique sequences.
Processing: 305047 
Sample 1 - 19447 reads in 2527 unique sequences.
Sample 1 - 19447 reads in 2419 unique sequences.
Processing: 305048 
Sample 1 - 4605 reads in 1035 unique sequences.
Sample 1 - 4605 reads in 1027 unique sequences.
Processing: 305049 
Sample 1 - 17641 reads in 3661 unique sequences.
Sample 1 - 17641 reads in 3319 unique sequences.
Processing: 305050 
Sample 1 - 21725 reads in 2411 unique sequences.
Sample 1 - 21725 reads in 2095 unique sequences.
Processing: 305051 
Sample 1 - 8890 reads in 1512 unique sequences.
Sample 1 - 8890 reads in 1610 unique sequences.
Processing: 305052 
Sample 1 - 12675 reads in 2465 unique sequences.
Sample 1 - 12675 reads in 2179 unique sequences.
Processing: 305053 
Sample 1 - 10890 reads in 1914 unique sequences.
Sample 1 - 10890 reads in 1661 unique sequences.
Processing: 305054 
Sample 1 - 12911 reads in 4312 unique sequences.
Sample 1 - 12911 reads in 3682 unique sequences.
Processing: 305056 
Sample 1 - 19733 reads in 2808 unique sequences.
Sample 1 - 19733 reads in 2576 unique sequences.
Processing: 305057 
Sample 1 - 12694 reads in 1639 unique sequences.
Sample 1 - 12694 reads in 1436 unique sequences.
Processing: 305058 
Sample 1 - 16816 reads in 1788 unique sequences.
Sample 1 - 16816 reads in 1522 unique sequences.
Processing: 305059 
Sample 1 - 16220 reads in 3448 unique sequences.
Sample 1 - 16220 reads in 3604 unique sequences.
Processing: 305060 
Sample 1 - 12869 reads in 1881 unique sequences.
Sample 1 - 12869 reads in 1508 unique sequences.
Processing: 305061 
Sample 1 - 15883 reads in 4207 unique sequences.
Sample 1 - 15883 reads in 3381 unique sequences.
Processing: 305062 
Sample 1 - 17427 reads in 4121 unique sequences.
Sample 1 - 17427 reads in 3334 unique sequences.
Processing: 305063 
Sample 1 - 16687 reads in 1888 unique sequences.
Sample 1 - 16687 reads in 1873 unique sequences.
Processing: 305064 
Sample 1 - 7962 reads in 1218 unique sequences.
Sample 1 - 7962 reads in 1040 unique sequences.
Processing: 305065 
Sample 1 - 14220 reads in 2052 unique sequences.
Sample 1 - 14220 reads in 1984 unique sequences.
Processing: 305066 
Sample 1 - 18357 reads in 2142 unique sequences.
Sample 1 - 18357 reads in 2153 unique sequences.
Processing: 305067 
Sample 1 - 17902 reads in 1740 unique sequences.
Sample 1 - 17902 reads in 1708 unique sequences.
Processing: 305068 
Sample 1 - 15676 reads in 2556 unique sequences.
Sample 1 - 15676 reads in 2234 unique sequences.
Processing: 305069 
Sample 1 - 15515 reads in 2991 unique sequences.
Sample 1 - 15515 reads in 2469 unique sequences.
Processing: 305070 
Sample 1 - 16520 reads in 3497 unique sequences.
Sample 1 - 16520 reads in 3128 unique sequences.
Processing: 305071 
Sample 1 - 14354 reads in 3205 unique sequences.
Sample 1 - 14354 reads in 2972 unique sequences.
Processing: 305072 
Sample 1 - 21998 reads in 4487 unique sequences.
Sample 1 - 21998 reads in 4250 unique sequences.
Processing: 305073 
Sample 1 - 6805 reads in 2423 unique sequences.
Sample 1 - 6805 reads in 2490 unique sequences.
Processing: 305074 
Sample 1 - 12507 reads in 1765 unique sequences.
Sample 1 - 12507 reads in 1403 unique sequences.
Processing: 305075 
Sample 1 - 16306 reads in 1730 unique sequences.
Sample 1 - 16306 reads in 1639 unique sequences.
Processing: 305076 
Sample 1 - 9980 reads in 1871 unique sequences.
Sample 1 - 9980 reads in 1629 unique sequences.
Processing: 305077 
Sample 1 - 13725 reads in 1651 unique sequences.
Sample 1 - 13725 reads in 1351 unique sequences.
Processing: 305078 
Sample 1 - 10257 reads in 1460 unique sequences.
Sample 1 - 10257 reads in 1895 unique sequences.
Processing: 305079 
Sample 1 - 20581 reads in 2062 unique sequences.
Sample 1 - 20581 reads in 2008 unique sequences.
Processing: 305080 
Sample 1 - 16190 reads in 3239 unique sequences.
Sample 1 - 16190 reads in 3254 unique sequences.
Processing: 305081 
Sample 1 - 18965 reads in 2763 unique sequences.
Sample 1 - 18965 reads in 3327 unique sequences.
Processing: 305082 
Sample 1 - 21372 reads in 5457 unique sequences.
Sample 1 - 21372 reads in 4787 unique sequences.
Processing: 305083 
Sample 1 - 14122 reads in 2047 unique sequences.
Sample 1 - 14122 reads in 1586 unique sequences.
Processing: 305084 
Sample 1 - 16937 reads in 1864 unique sequences.
Sample 1 - 16937 reads in 1318 unique sequences.
Processing: 305085 
Sample 1 - 3329 reads in 1840 unique sequences.
Sample 1 - 3329 reads in 1959 unique sequences.
Processing: 305086 
Sample 1 - 18411 reads in 2248 unique sequences.
Sample 1 - 18411 reads in 2091 unique sequences.
Processing: 305087 
Sample 1 - 18328 reads in 2261 unique sequences.
Sample 1 - 18328 reads in 1707 unique sequences.
Processing: 305088 
Sample 1 - 15051 reads in 1736 unique sequences.
Sample 1 - 15051 reads in 1602 unique sequences.
Processing: 305089 
Sample 1 - 22257 reads in 2560 unique sequences.
Sample 1 - 22257 reads in 2467 unique sequences.
Processing: 305090 
Sample 1 - 815 reads in 417 unique sequences.
Sample 1 - 815 reads in 410 unique sequences.
Processing: 305091 
Sample 1 - 18885 reads in 2364 unique sequences.
Sample 1 - 18885 reads in 1907 unique sequences.
Processing: 305092 
Sample 1 - 21374 reads in 5466 unique sequences.
Sample 1 - 21374 reads in 4237 unique sequences.
Processing: 305093 
Sample 1 - 20498 reads in 2577 unique sequences.
Sample 1 - 20498 reads in 2442 unique sequences.
Processing: 305094 
Sample 1 - 2629 reads in 684 unique sequences.
Sample 1 - 2629 reads in 671 unique sequences.
Processing: 305095 
Sample 1 - 3315 reads in 986 unique sequences.
Sample 1 - 3315 reads in 1094 unique sequences.
Processing: 305096 
Sample 1 - 13718 reads in 1801 unique sequences.
Sample 1 - 13718 reads in 1964 unique sequences.
Processing: 305097 
Sample 1 - 18857 reads in 1639 unique sequences.
Sample 1 - 18857 reads in 1495 unique sequences.
Processing: 305098 
Sample 1 - 13082 reads in 3119 unique sequences.
Sample 1 - 13082 reads in 3239 unique sequences.
Processing: 305099 
Sample 1 - 15432 reads in 1628 unique sequences.
Sample 1 - 15432 reads in 1319 unique sequences.
Processing: 305100 
Sample 1 - 5650 reads in 1487 unique sequences.
Sample 1 - 5650 reads in 1362 unique sequences.
Processing: 305101 
Sample 1 - 12952 reads in 1224 unique sequences.
Sample 1 - 12952 reads in 1042 unique sequences.
Processing: 305102 
Sample 1 - 15731 reads in 1888 unique sequences.
Sample 1 - 15731 reads in 2115 unique sequences.
Processing: 305103 
Sample 1 - 13258 reads in 1729 unique sequences.
Sample 1 - 13258 reads in 1541 unique sequences.
Processing: 305104 
Sample 1 - 15497 reads in 2490 unique sequences.
Sample 1 - 15497 reads in 2409 unique sequences.
Processing: 305105 
Sample 1 - 19793 reads in 2039 unique sequences.
Sample 1 - 19793 reads in 2136 unique sequences.
Processing: 305106 
Sample 1 - 14035 reads in 1438 unique sequences.
Sample 1 - 14035 reads in 1152 unique sequences.
Processing: 305107 
Sample 1 - 52 reads in 31 unique sequences.
Sample 1 - 52 reads in 28 unique sequences.
Processing: 305108 
Sample 1 - 17937 reads in 3469 unique sequences.
Sample 1 - 17937 reads in 3121 unique sequences.
Processing: 305109 
Sample 1 - 12873 reads in 2667 unique sequences.
Sample 1 - 12873 reads in 2551 unique sequences.
Processing: 305110 
Sample 1 - 17 reads in 14 unique sequences.
Sample 1 - 17 reads in 14 unique sequences.
Processing: 305111 
Sample 1 - 16349 reads in 1770 unique sequences.
Sample 1 - 16349 reads in 1578 unique sequences.
Processing: 305112 
Sample 1 - 54 reads in 38 unique sequences.
Sample 1 - 54 reads in 41 unique sequences.
Processing: 305113 
Sample 1 - 15402 reads in 1792 unique sequences.
Sample 1 - 15402 reads in 1614 unique sequences.
Processing: 305114 
Sample 1 - 5 reads in 4 unique sequences.
Sample 1 - 5 reads in 5 unique sequences.
Processing: 305115 
Sample 1 - 12918 reads in 2785 unique sequences.
Sample 1 - 12918 reads in 2699 unique sequences.
Processing: 305116 
Sample 1 - 19471 reads in 2605 unique sequences.
Sample 1 - 19471 reads in 1808 unique sequences.
Processing: 305117 
Sample 1 - 18656 reads in 2394 unique sequences.
Sample 1 - 18656 reads in 2302 unique sequences.
Processing: PBS1 
Sample 1 - 233 reads in 121 unique sequences.
Sample 1 - 233 reads in 136 unique sequences.
Processing: PBS2 
Sample 1 - 98 reads in 65 unique sequences.
Sample 1 - 98 reads in 60 unique sequences.
Processing: PBS3 
Sample 1 - 13 reads in 12 unique sequences.
Sample 1 - 13 reads in 12 unique sequences.
Processing: PBS4 
Sample 1 - 92 reads in 29 unique sequences.
Sample 1 - 92 reads in 30 unique sequences.
> 
>   rm(derepF); rm(derepR)
> 
>   ## Make sequence abundance table 
>   seqtab <- makeSequenceTable(mergers)
The sequences being tabled vary in length.
>   saveRDS(seqtab, "dada2_abundance_table.rds")
> 
>   getN <- function(x) sum(getUniques(x))
>   ## track <- cbind(out, rowSums(seqtab))
>   v<-rowSums(seqtab)
>   v0<-numeric(nrow(out))
>   track<-cbind(out, v0)
>   rownames(track)<-gsub("_R1_tc.fastq","",rownames(track))
>   track[names(v),3]<-v
>   colnames(track) <- c("input", "filtered", "merged")
>   write.table(track, "dada2_part1_stats.txt", quote=FALSE, append=FALSE, sep=	, row.names=TRUE, col.names=TRUE)
> path<-getwd()
>
> ## list all of the files matching the pattern
> tables<-list.files(path, pattern="-dada2_abundance_table.rds", full.names=TRUE)
> stats<-list.files(path, pattern="-dada2_part1_stats.txt", full.names=TRUE)
>
> ## get the run names using splitstring on the tables where - exists
> sample.names <- sapply(strsplit(basename(tables), "-"), `[`, 1)
> ##sample.names
> ##names(tables) <- sample.names
>
> runs <- vector("list", length(sample.names))
> names(runs) <- sample.names
> for(run in tables) {
+   cat("Reading in:", run, "
+ ")
+   runs[[run]] <- readRDS(run)
+ }
Reading in: /local/groupshare/ravel/jain/2_Step_MiSeq_Meno/2_Step_MiSeq_Meno_RAV454_461-dada2_abundance_table.rds
>
> runstats <- vector("list", length(sample.names))
> names(runstats) <- sample.names
> for(run in stats) {
+   cat("Reading in:", run, "
+ ")
+   runstats[[run]] <- read.delim(run, )
+ }
Reading in: /local/groupshare/ravel/jain/2_Step_MiSeq_Meno/2_Step_MiSeq_Meno_RAV454_461-dada2_part1_stats.txt
>
> unqs <- unique(c(sapply(runs, colnames), recursive=TRUE))
> n<-sum(unlist(lapply(X=runs, FUN = nrow)))
> st <- matrix(0L, nrow=n, ncol=length(unqs))
> rownames(st) <- c(sapply(runs, rownames), recursive=TRUE)
> colnames(st) <- unqs
> for(sti in runs) {
+   st[rownames(sti), colnames(sti)] <- sti
+ }
> st <- st[,order(colSums(st), decreasing=TRUE)]
>
> ##st.all<-mergeSequenceTables(runs)
> # Remove chimeras
> seqtab <- removeBimeraDenovo(st, method="consensus", multithread=TRUE)
> # Assign taxonomy
> silva <- assignTaxonomy(seqtab, "/home/jholm/bin/silva_nr_v128_train_set.fa.gz", multithread=TRUE)
> # Write to disk
> saveRDS(seqtab, "all_runs_dada2_abundance_table.rds") # CHANGE ME to where you want sequence table saved
> write.csv(seqtab, "all_runs_dada2_abundance_table.csv", quote=FALSE)
> write.csv(silva, "silva_classification.csv", quote=FALSE)
>
> fc = file("all_runs_dada2_ASV.fasta")
> fltp = character()
> for( i in 1:ncol(seqtab))
+ {
+   fltp <- append(fltp, paste0(">", colnames(seqtab)[i]))
+   fltp <- append(fltp, colnames(seqtab)[i])
+ }
> writeLines(fltp, fc)
> rm(fltp)
> close(fc)
>
> track<-as.matrix(rowSums(seqtab))
> colnames(track) <- c("nonchimeric")
> write.table(track, "dada2_part2_stats.txt", quote=FALSE, append=FALSE, sep="  ", row.names=TRUE, col.names=TRUE)
