## Supplemental File 4 for "Ultra-high throughput multiplexing and sequencing of >500 bp amplicon regions on the Illumina HiSeq 2500 platform"

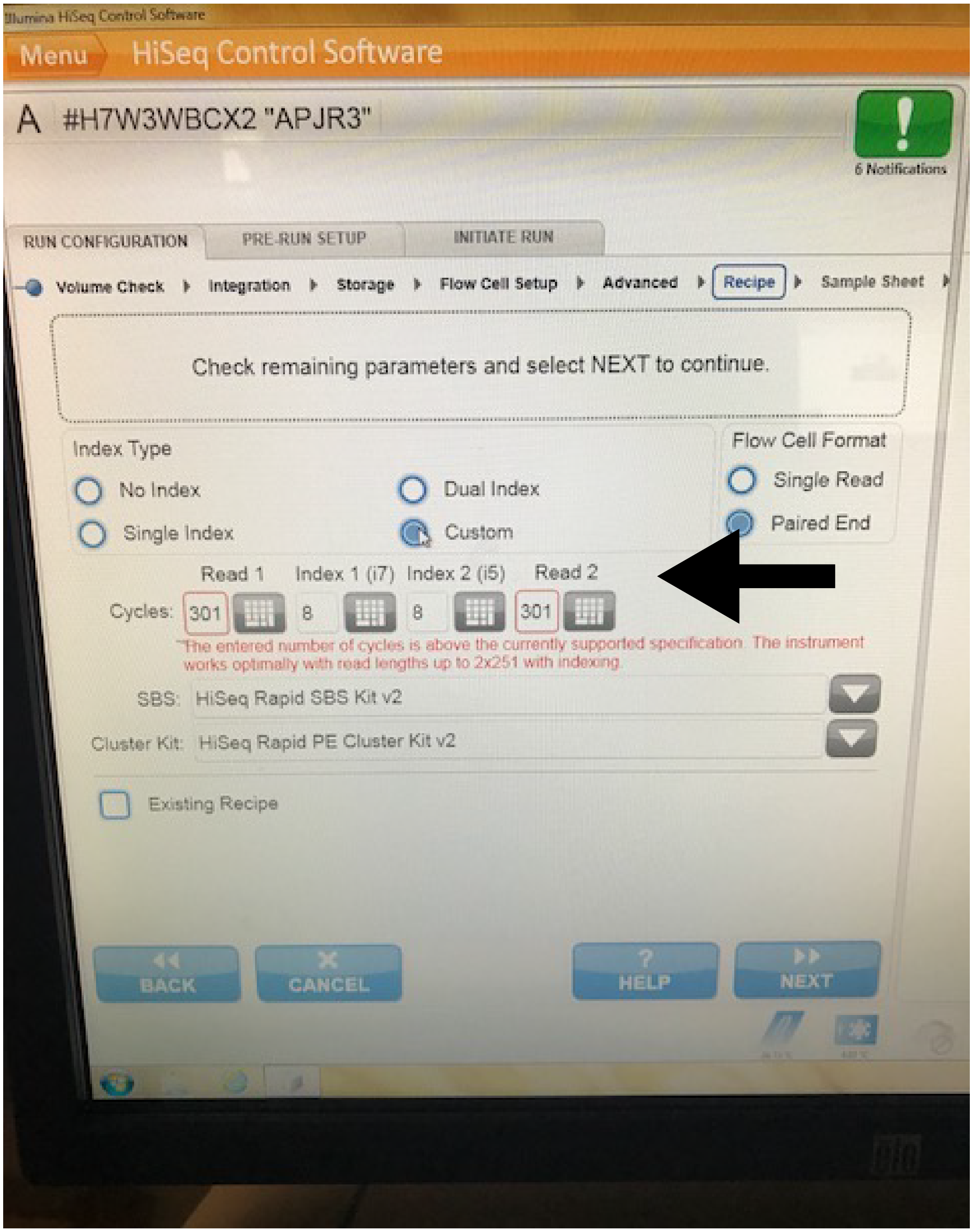


Supplementary File 4. HiSeq Control Software configuration for obtaining 2 x 300 bp paired-end reads using the Illumina HiSeq 2500.
