## Supplemental File 5 for "Ultra-high throughput multiplexing and sequencing of >500 bp amplicon regions on the Illumina HiSeq 2500 platform"

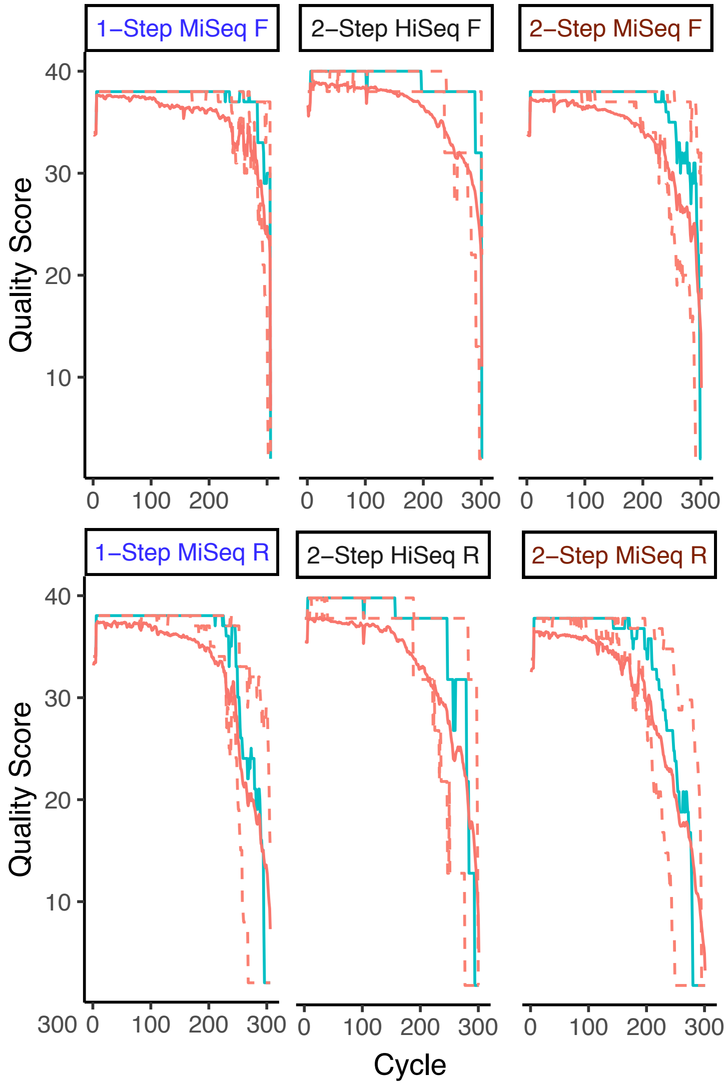


Supplemental File 6. Per cycle quality score summaries for vaginal samples prepared and sequenced using the 1-Step PCR method and Illumina MiSeq (n=49 samples), 2-Step PCR method and Illumina MiSeq (n=83 samples), and 2-Step PCR method and Illumina HiSeq (n=83 samples), Shown for each cycle are the mean quality score (green line), the median quality score (solid orange line), the quartiles of the quality score distribution (dotted orange lines).
