## Supplemental File 6 for "Ultra-high throughput multiplexing and sequencing of >500 bp amplicon regions on the Illumina HiSeq 2500 platform"

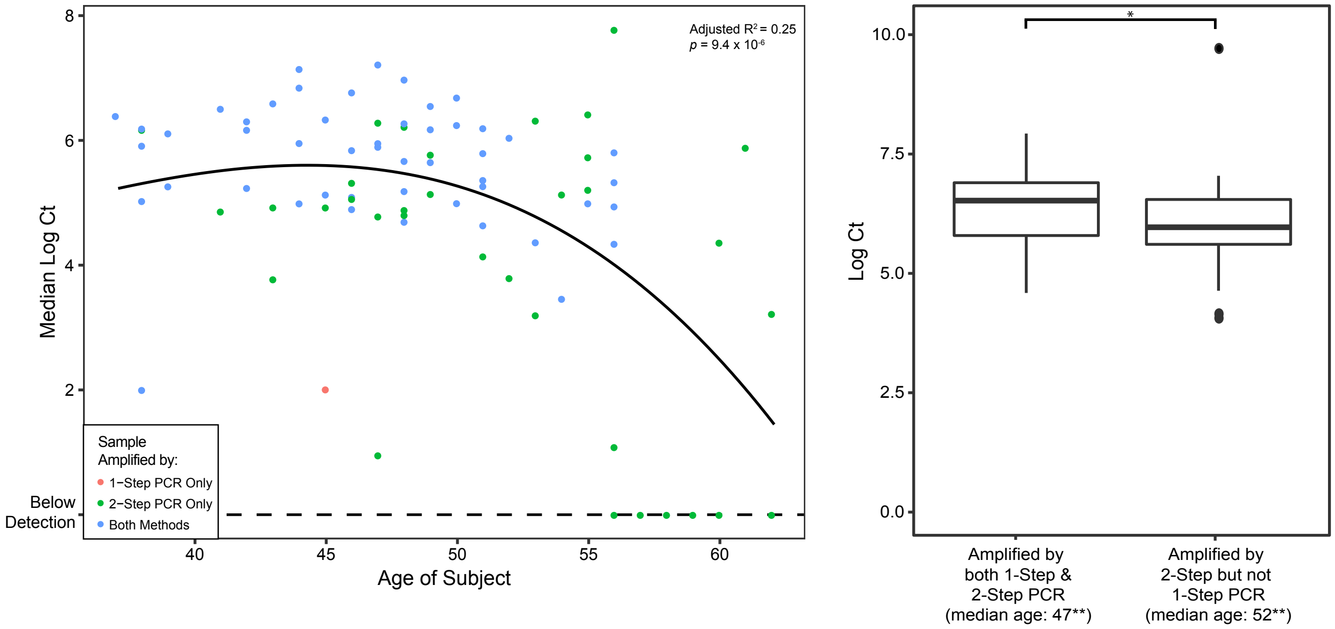


Supplemental Figure 7. Pan-bacterial qPCR determination of the number of 16S rRNA gene targets in vaginal samples from women of varying ages (left). Vaginal samples which were amplified by 2-Step PCR but not 1-Step PCR had significantly-lower abolute abundances of 16S rRNA gene targets (U = 790.5, p = 0.027), and subjects from which samples came were siginficantly older (U = 484.5, p = 0.001).
