## Supplemental File 7 for "Ultra-high throughput multiplexing and sequencing of >500 bp amplicon regions on the Illumina HiSeq 2500 platform"

1 Results
2 Materials and Methods
3 QC Data
4 References

### ZymoBIOMICS® Service Report

*Zymo Research Corporation*

### 1 Results

**Click on the links below to view the results from each analysis:**

**StandardQC.Bac16Sv34 (./StandardQC.Bac16Sv34/results.html)**

### 2 Materials and Methods

The samples were processed and analyzed with the ZymoBIOMICS® Service: Targeted Metagenomic Sequencing (Zymo Research, Irvine, CA).

**DNA Extraction:** One of three different DNA extraction kits was used depending on the sample type and sample volume. In most cases, the ZymoBIOMICS® DNA Miniprep Kit (Zymo Research, Irvine, CA) was used. For low biomass samples, such as skin swabs, the ZymoBIOMICS® DNA Microprep Kit (Zymo Research, Irvine, CA) was used as it permits for a lower elution volume, resulting in more concentrated DNA samples. For a large sample volume, the ZymoBIOMICS®-96 MagBead DNA Kit (Zymo Research, Irvine, CA) was used to extract DNA using an automated platform.

**Targeted Library Preparation:** Bacterial 16S ribosomal RNA gene targeted sequencing was performed using the Quick-16S™ NGS Library Prep Kit (Zymo Research, Irvine, CA). The bacterial 16S primers amplified the V1-V2 or V3-V4 region of the 16S rRNA gene. These primers have been custom-designed by Zymo Research to provide the best coverage of the 16S gene while maintaining high sensitivity. Fungal ITS gene targeted sequencing was performed using the Quick-16S™ NGS Library Prep Kit with custom ITS2 primers substituted for 16S primers.

The sequencing library was prepared using an innovative library preparation process in which PCR reactions were performed in real-time PCR machines to control cycles and therefore prevent limit PCR chimera formation. The final PCR products are were quantified with qPCR fluorescence readings and pooled together based on equal molarity. The final pooled library was cleaned up with the Select-a-Size DNA Clean & Concentrator™ (Zymo Research, Irvine, CA), then quantified with TapeStation® and Qubit®.

**Sequencing:** The final library was sequenced on Illumina® MiSeq™ with a v3 reagent kit (600 cycles). The sequencing was performed with >10% PhiX spike-in.

**Bioinformatics Analysis:** Unique amplicon sequences were inferred from raw reads using the Dada2 pipeline (Callahan et al, 2016). Chimeric sequences were also removed with the Dada2 pipeline. Typically, two bioinformatics reports were generated that differ in the database used.In both, taxonomy assignment was performed using Uclust from Qiime v.1.9.1. In the report named with *.zymo*, taxonomy was assigned with the Zymo Research Database, a 16S database that is internally designed and curated, as reference. In the report named with *.greengene*, taxonomy was assigned with the Greengenes 16S database as reference. Composition visualization, alpha-diversity, and beta-diversity analyses were performed with Qiime v.1.9.1 (Caporaso et al., 2010). If applicable, taxonomy that have significant abundance among different groups were identified by LEfSe (Segata et al., 2011) using default settings. Other analyses such as heatmaps, Taxa2SV_deomposer, and PCoA plots were performed with internal scripts.

### 3 QC Data

Zymo Research's microbiomics workflows include sufficient quality controls. The ZymoBIOMICS® Microbial Community Standards (both cellular standard and DNA standard) were used as positive controls for each run. Negative controls (e.g. blank extraction control, blank library preparation control) were included to assess the level of bioburden carried by the wet-lab process. The barplot below shows the microbial composition of the ZymoBIOMICS® Microbial Community Standards measured in this project. The theoretical composition of the standard can be accessed with the link given below the figure.

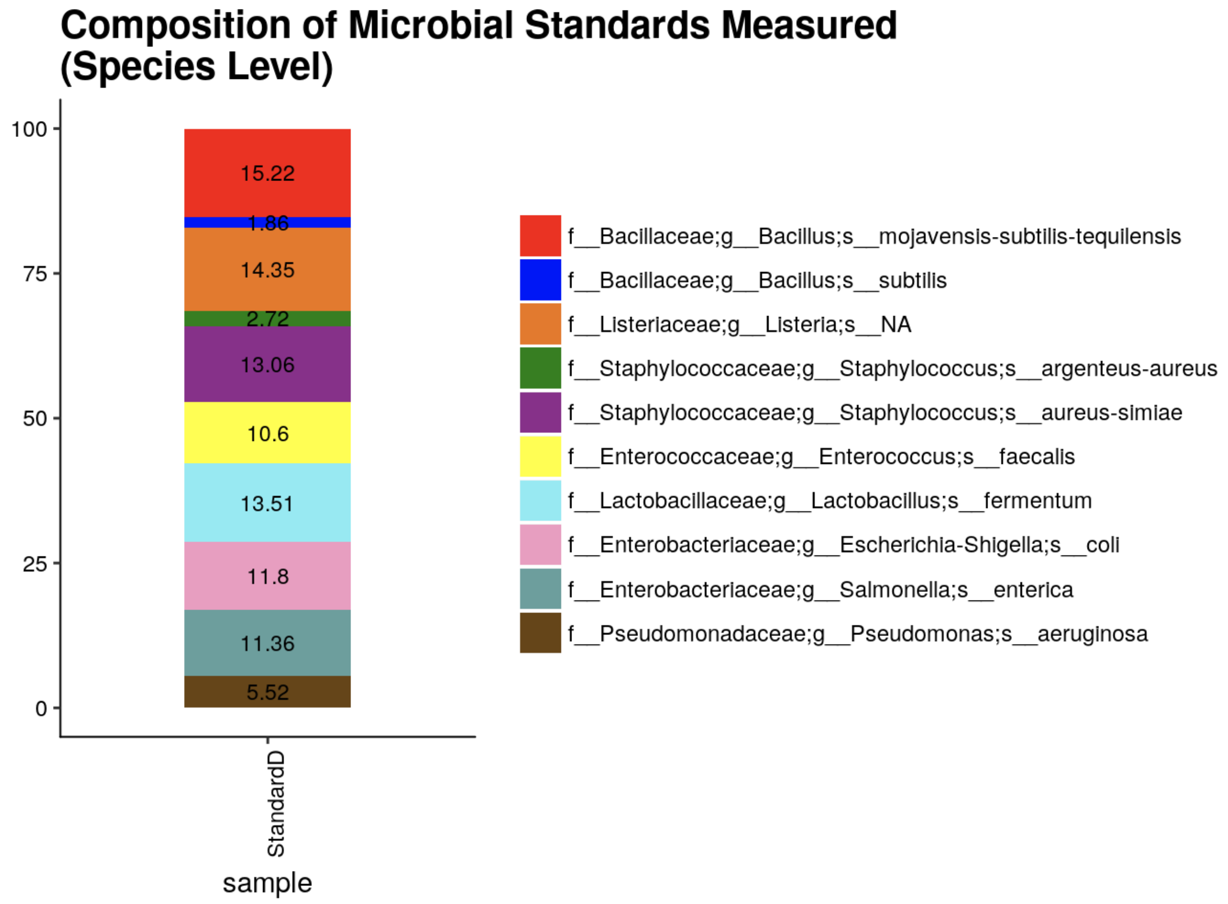

**Theoretical Composition of the ZymoBIOMICS® Microbial Community Standard.**

 (mailto:)

www.zymoresearch.com (https://www.zymoresearch.com/other-services?service=microbiomics) Toll Free: (888) 882-9682
