## Supplemental File 8 for "Ultra-high throughput multiplexing and sequencing of >500 bp amplicon regions on the Illumina HiSeq 2500 platform"

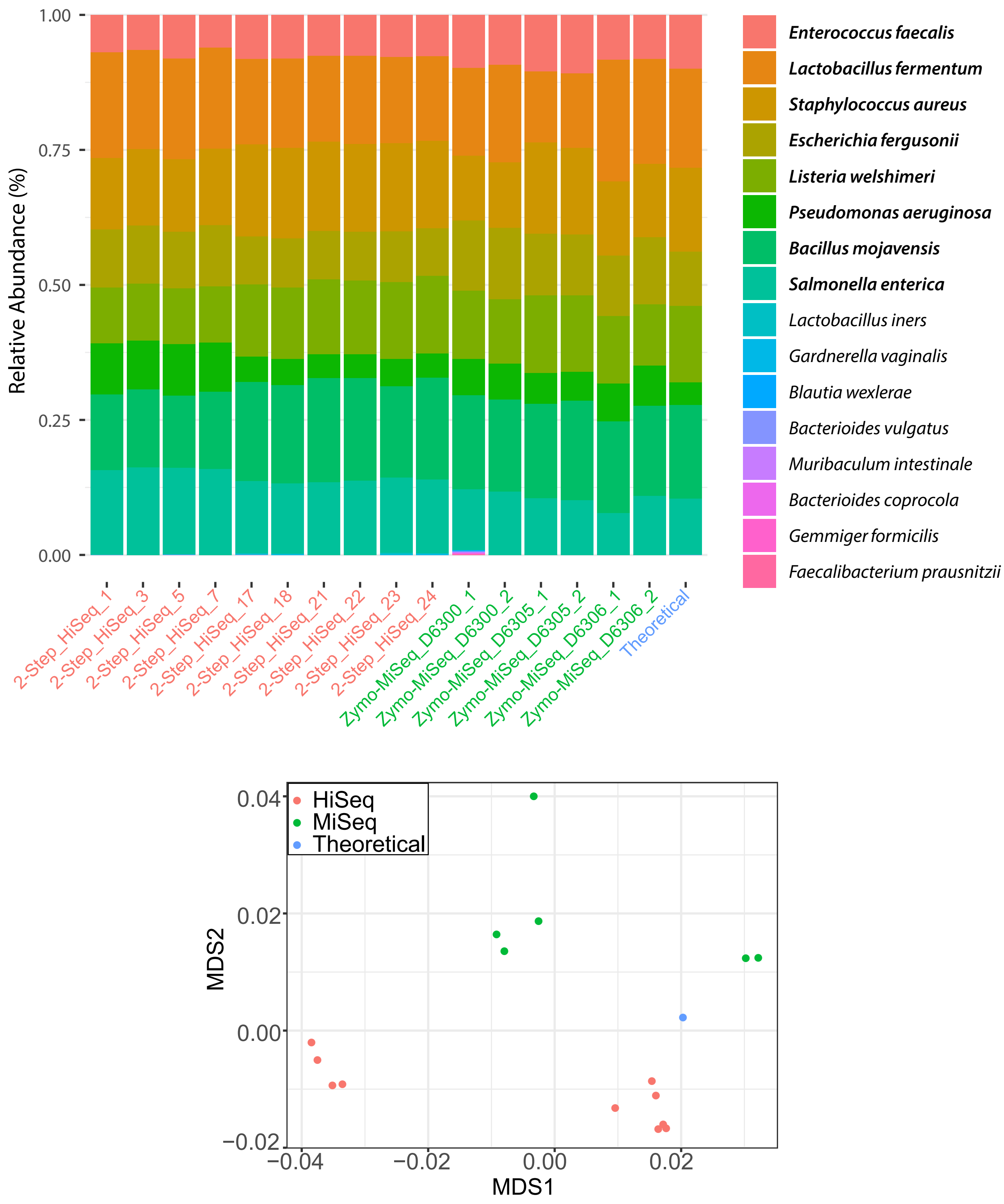


Supplementary File 9. Top: Microbial compositions of the Zymobiomics Microbial DNA Standard prepared and sequenced using the 2-Step library prepartion method described in this paper and the Illumina HiSeq platform (2-Step_HiSeq), or Zymo-prepared and Illumina MiSeq- sequenced (Zymo-MiSeq) are statistically similar when compared to the theoretical values reported by Zymo. Taxa in bold compose the mock communities. Bottom: Multidimensional scaling plot of Jensen-Shannon distances of the same samples as top.
