## Supplementary figures and images for "Ultra-high throughput multiplexing and sequencing of >500 bp amplicon regions on the Illumina HiSeq 2500 platform"

### Supplemental File 9

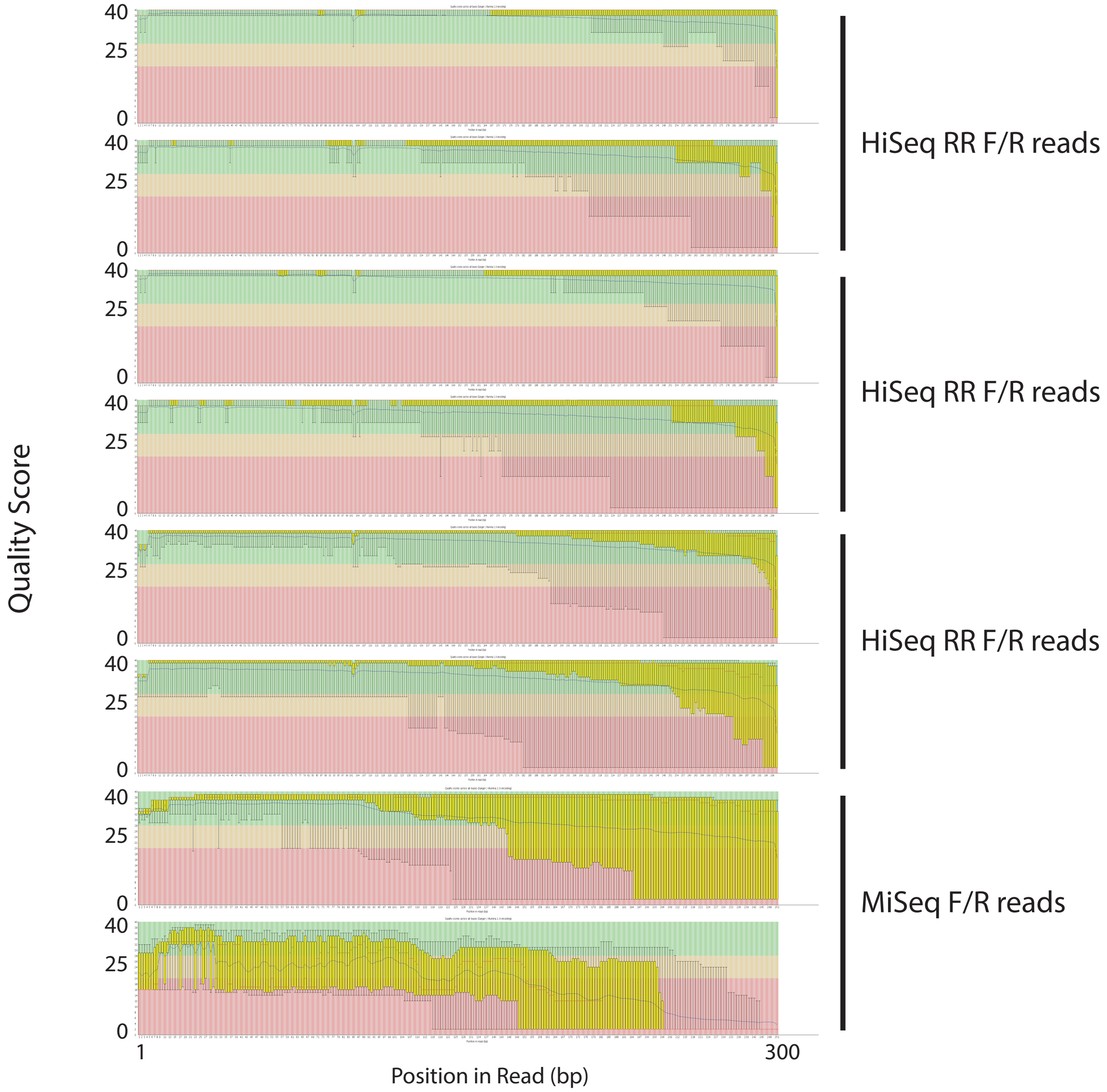


Supplementary File 10. FastQC output from 3 HiSeq Rapid Run (HiSeq RR) and MiSeq sequencing runs.
